# Simplified post-assembly plasmid library amplification for increased transformation yields in *E. coli* and *S. cerevisiae*

**DOI:** 10.1101/2023.11.02.565362

**Authors:** Thomas Fryer, Darian S. Wolff, Max D. Overath, Elena Schäfer, Andreas H. Laustsen, Timothy P. Jenkins, Carsten Andersen

**Affiliations:** Department of Biotechnology and Biomedicine, Technical University of Denmark, Kongens Lyngby, Denmark; Department of Molecular Discovery, R&D, Novozymes A/S; Department of Biochemistry, University of Cambridge

**Keywords:** Directed evolution, DNA library preparation, Rolling Circle Amplification, transformation efficiency, *in vivo* homology directed assembly

## Abstract

Many biological disciplines rely upon the transformation of host cells with heterologous DNA, the limited efficiency of which can significantly hinder a researcher’s work. Directed evolution in particular typically requires the screening of large (thousand-billion member) libraries to identify sequences of interest, and the creation of these libraries at large enough scales to overcome transformation inefficiencies is a cost-and time-intensive process. We simplify this process by using Rolling Circle Amplification (RCA) to amplify *in vitro* plasmid DNA assembly reactions, followed by facile resolution of the concatomeric products to monomers through treatment with specific endonucleases and subsequent efficient transformation of the linear DNA products into host cells for *in vivo* circularisation. We demonstrate that use of a nicking endonuclease to generate homologous single-stranded ends increases the efficiency of *E. coli* chemical transformation versus both linear DNA with double-stranded homologous ends, and circular golden-gate assembly products, whilst use of a restriction endonuclease to generate linear DNA with double-stranded homologous ends increases the efficiency of chemical and electrotransformation of *S. cerevisiae*. Importantly, we also optimise the process such that both RCA and endonuclease treatment occur efficiently in the same buffer, streamlining the workflow and reducing product loss through purification steps. We expect our approach to have utility beyond directed evolution in *E. coli* and *S. cerevisiae*, to areas such as genome engineering and the manipulation of alternative organisms with even poorer transformation efficiencies.

## Introduction

The manipulation of DNA and its subsequent insertion into a desired host cell is a bedrock of modern biotechnology. This ability has advanced both basic science (e.g., the human genome project, metagenomics, gene and protein function assays), as well as applied science (e.g., the directed evolution of proteins for a desired functionality, and the production of proteins and small molecules via fermentation and biocatalytic processes). To achieve DNA transfer into microorganisms there are three main routes: conjugation, transduction, and transformation, in which DNA is transferred cell-to-cell, virus-to-cell, or extracellular DNA-to-cell, respectively. Transformation is the most routinely used DNA transfer mechanism in laboratory settings due to its relative ease (eschewing any need for viral packaging reactions^1^ or co-culturing of conjugative cells^2^) and historical precedent, in which protocols have been established for many different cell types across all kingdoms of life. In particular, many protocols have been established for two of the most common organisms used in biotechnology, *E. coli*^3^ and *S. cerevisiae*^4^, that are broadly separable into either chemical competent or electrocompetent methods. Briefly, chemical competent approaches typically rely on washing in mixtures of salts followed by subsequent incubation with exogenous DNA that is induced to pass through the cell membrane by a heat-shock step, whilst electrocompetent approaches wash cells in hypertonic solutions to remove any ions and then use exposure to electrical current to drive exogenous DNA through the cellular membrane^5^. Despite the vast number of reported protocols, transformation is still limited by relatively poor efficiencies, with typically <1% of cells being transformed by exogenous DNA. Particularly in the field of directed evolution, it is desirable to make libraries that are as large and high quality as possible, so that the greatest amount of sequence space can be explored in the search for a desired trait^6^. Limited transformation efficiencies greatly hinder this work, and are compounded by the discrepancies between advertised transformation efficiencies (measured using picogram quantities of supercoiled plasmid DNA) and those achieved during library preparations (using nanogram-microgram quantities of relaxed circular DNA) often resulting in 100-1000-fold loss in efficiency.

In order to overcome limited transformation efficiencies, researchers typically scale-up the process by transforming multiple aliquots of cells using as much DNA as possible. Whilst multiple aliquots of cells can either be purchased (at significant cost) or prepared in-house with relative ease, the preparation of large quantities (10-100 micrograms) of library DNA is more difficult. To achieve large quantities of assembled library DNA, one can either scale-up the assembly reactions or amplify the assembled product in a way that it is suitable for transformation into host cells. There are now many different ways to assemble libraries beyond the traditional restriction-ligation approach, many of which offer the benefit of ‘scarless’ assembly, *i*.*e*., the absence of any undesired sequence in the final product. The most common scarless approaches are Gibson assembly^7^ and golden-gate assembly^8^ encountered in the form of commercial kits, which cause the scale-up of library assemblies to quickly become economically infeasible for many labs (overviewed nicely in Xia et al.^9^). Whilst much work has gone into non-commercial derivatives^10^ or alternatives^9,11^, these are often non-robust or complex to establish. Perhaps the most established non-commercial scarless cloning approach utilises the native ability of multiple cell types to *in vivo* repair linear DNA into circular plasmid DNA. Whilst it is widely known that *S. cerevisiae* is transformable by linear DNA with homologous ends^12^, it is often overlooked that *E. coli* possesses the same ability^13^ (albeit likely occurring through a different non-recombination-based pathway). As these approaches unite the plasmid assembly and transformation step in a cost-free manner, they are particularly scalable and therefore suitable for large library generation. Thus, a method to generate large quantities of DNA in a suitable format (*i*.*e*. linear DNA) is required. Much work on amplifying library DNA from small-scale assemblies has already been carried out, typically focussing on Rolling Circle Amplification (RCA) due to its ability to massively amplify circular DNA in an isothermal manner. The product of RCA (highly-branched concatomers) is however not directly suitable for transformation and must be resolved into monomeric units, achieved in the literature through digestion with restriction enzymes and subsequent large-volume low concentration religation^14^ or Cre recombinase activity^15^. Whilst both approaches are successful, they are either cumbersome or require non-standard reagents. A recent elegant development utilises a one-pot golden-gate RCA reaction, which simplifies the generation of large quantities of cloned DNA. However, the cloned DNA is generated only in a final circular format and without a quantification of its impact on the generation of large libraries^16^. To best deliver a method that can easily generate large libraries in multiple different cell types, we thus sought to combine features from multiple different approaches. Notably, we utilised RCA to amplify a small-scale golden-gate assembly reaction ∼10,000 fold generating ∼100 ug of product, followed by nicking endonuclease (Nb. BbvCI) treatment (*E. coli* workflow) to generate linear DNA with single-stranded homologous ends or blunt-ended restriction endonuclease (FspI) treatment (*S. cerevisiae* workflow) to generate linear DNA with double-stranded homologous ends. Subsequently, these linear DNA molecules can be efficiently transformed into the appropriate cell types.

## Results

### Design of post-assembly plasmid amplification

Focusing firstly on E. coli, we designed our approach around three key techniques: 1) RCA of small-scale library assemblies to generate ∼100 ug of DNA, 2) nicking endonuclease treatment to resolve the RCA product into linear monomeric units, and 3) *in vivo* assembly of linear DNA by *E. coli* to generate circular plasmids (Figure 1A). Nicking endonuclease treatment was chosen based on observations from Xia et al.^9^ that generation of single-stranded homologous ends are sufficient to recapitulate the cloning and transformation efficiencies of more complex Gibson assemblies, and are more efficient than the transformation of linear DNA with double-stranded homologous ends seen in García-Nafría et al.^13^. Additionally, use of nicking endonucleases for cloning purposes has been successfully developed by Wang et al.^17^. Our test plasmid was thus modified to include a ‘nickase cassette’, in which two Nb. BbvCI sites were added on the top and bottom strand of the plasmid respectively, such that upon nickase treatment a) the concatomeric RCA product will be resolved to monomers and b) the linear monomers will possess single-stranded 5’ and 3’ overhangs that are homologous to one another.

**Figure 1.**
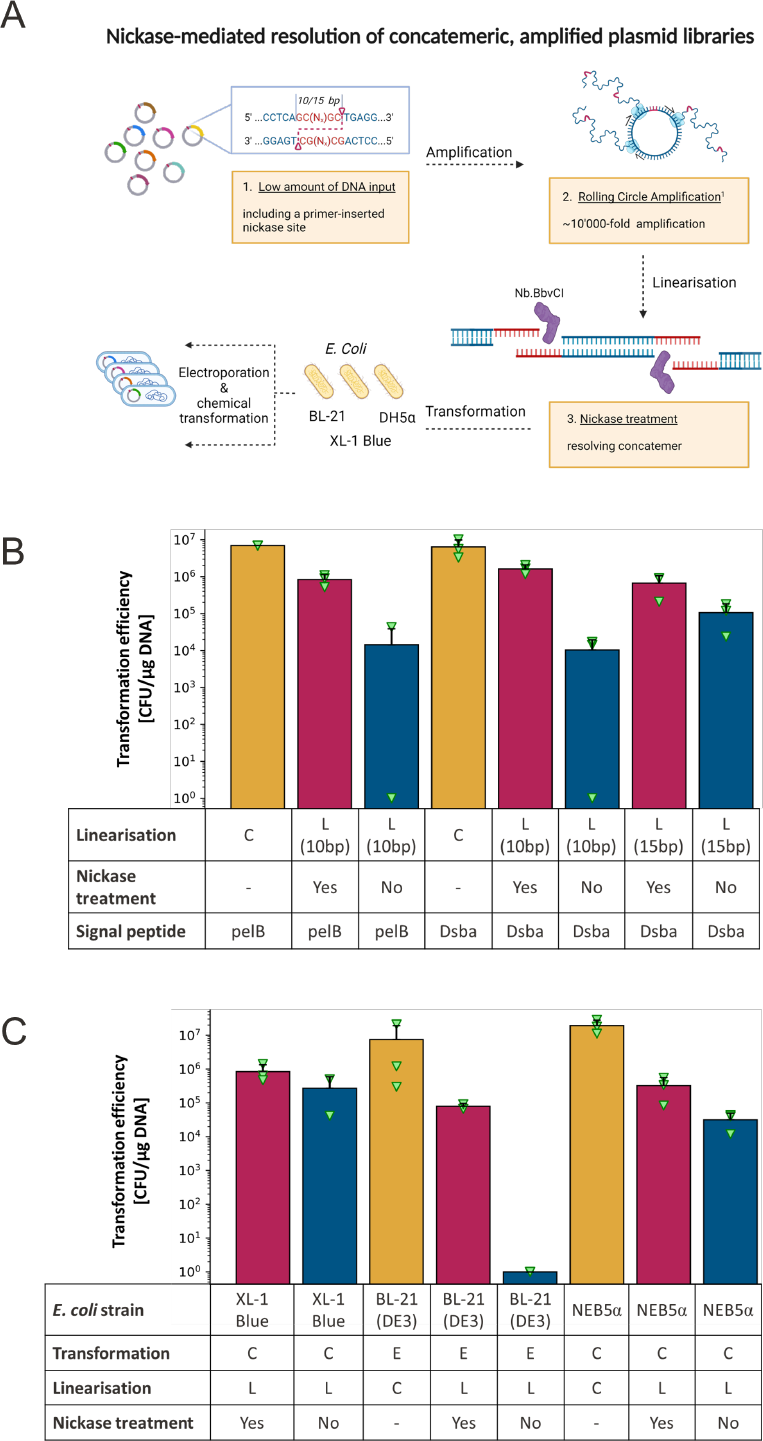
Testing the influence of sticky or blunt DNA overhangs while using supercoiled DNA as control for transformability of different *E. coli* cell strains. **(a)** Optimised post-assembly library amplification for transformation into a variety of *E. coli* strains (*e*.*g*. XL-1 Blue, TG1, BL21 (DE3), and NEB5α). For simplicity RCA is displayed without further branching of the first layer of elongated product.(**b**) Comparison of transformation efficiencies in the same *E. coli* strain (XL-1 Blue) and the VHH-encoding plasmid (‘PF-Nbb102-CAM’) across the different (1) formats (circular (orange), single-stranded homology site (red), double-stranded homology site (blue)), (2) a varying base pair length of the homology site (10 and 15 bp), and (3) encoding for a different signal-peptide (pelB and Dsba). The abbreviations ‘C’ and ‘L’ in the row ‘linearisation’ refer to plasmid formats ‘circular/supercoiled’ and ‘linear’, respectively. (**c**) Assessing transformation efficiency of the same VHH plasmid (‘PF-Nbb102-CAM’) in circular (orange) and linear format across different *E. coli* strains (XL-1 Blue, BL21 (DE3), and NEB5α). The linear format is further divided into plasmids with single-stranded homology sites (nickase-treated) (red) and double-stranded homology sites (non-nickase treated) (blue). Importantly, no rolling circle amplification was conducted prior to the experiment and all linear DNA products are from PCR amplification with appropriate primers to add the nickase sites. Furthermore, the abbreviations ‘E’ and ‘C’ in the row ‘transformation’ stand for the transformation method ‘electroporation’ and ‘chemical transformation’, which are described in detail in the method section. Data is displayed as the mean of triplicates (with individual data points displayed with green triangles).Error bars represent standard deviation of replicates nand face only upwards for simplicity.

### Validation of single-strand overhangs as enhancers of transformation efficiency

As a first test of our approach, we investigated the effect of creating single-stranded homologous ends on transformation efficiency across multiple cell types and homology lengths. Initially, two different homology lengths were tested (10 bases and 15 bases) by adding the appropriate sequences through PCR amplification of the template plasmid (a phagemid for a VHH phage display library, with either the pelB or DsbA signal peptides for periplasmic export of the VHH-p3 fusion protein). When subjecting the linear DNA products to nickase treatment, a substantial boost in transformation efficiency (10-100 times higher) was observed compared to linear DNA with double-stranded homologous ends for both homology lengths. Notably, the 10-base homology length exhibited a slightly superior transformation efficiency relative to the 15-base homology length in the context of the DsbA signal peptide (see Figure 1B). As such, we proceeded with the 10 base homology length and additionally tested the transformation efficiency of a pelB signal peptide containing phagemid in comparison to the DsbA containing phagemids. Little-to-no effect was seen, thus we proceeded with the more standard pelB signal peptide in subsequent experiments. We then explored the transformation of different cell types relevant to directed evolution (XL-1 blue for phage display, BL21-DE3 for protein expression, and NEB5α for cloning) using nickase-treated linear DNA, and observed the same effects as before, *i*.*e*., that single-stranded homologous ends enhanced transformation efficiency when compared to double-stranded homologous ends across all cell types (Figure 1C).

### Nickase-mediated resolution of concatomeric RCA products

After confirming that nickase treatment was beneficial, we investigated the best conditions for RCA of plasmids focussing on both amplification yield and specificity based on literature protocols^18,19^. Importantly, the plasmid of interest now contained the 10 bp nickase cassette that resulted from transformation of the linear DNA products in Figure 1. This investigation uncovered that NEB Phi29 coupled with RNA or DNA (phosphorothioate-protected) random hexamers yielded the most product (>10,000 fold amplification) (Figure 2A). We also observed that amplification yield increased linearly with reduced input template DNA concentration (Figure 2B), suggesting that a saturating quantity of product DNA is reached independent of input template concentration. Next, we sought to confirm that treatment with the nickase would resolve the concatomeric DNA product into monomeric units through agarose gel electrophoresis (Figure 2C). Successful resolution was observed, thus, we investigated whether the RCA-nickase workflow could be simplified, such that minimal buffer exchange or purification would be required. This approach identified the use of 1x NEB rCutSmart buffer with 0.2 mg/mL NEB rAlbumin as the most efficient, conveniently enabling both RCA and nickase treatment without purification or buffer exchange.

**Figure 2.**
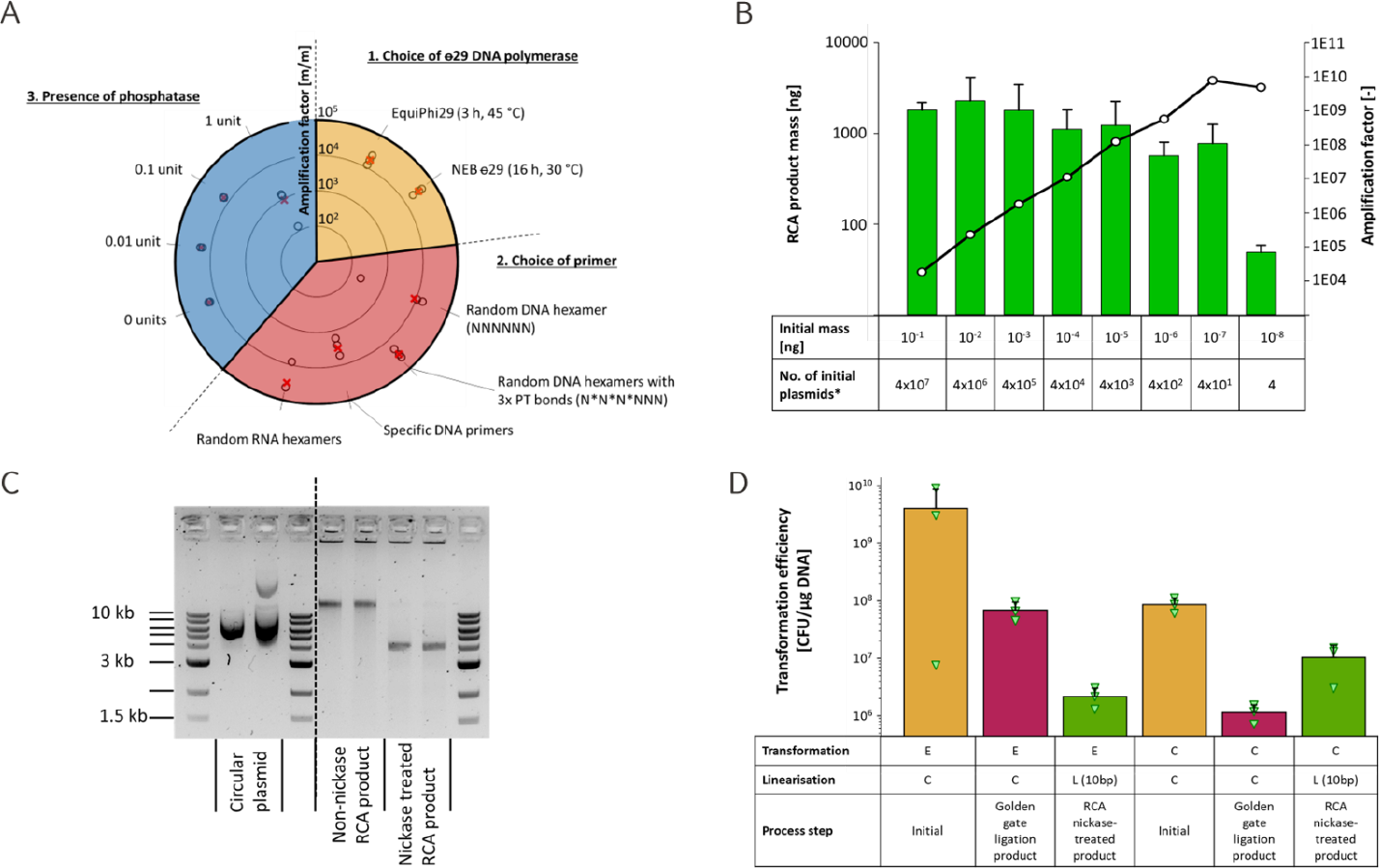
Robust and simplified library amplification using RCA. (**a**) Seeking to optimise conditions for rolling circle amplification, several parameters were considered and evaluated when test-amplifying the plasmid ‘PF-Nbb102-CAM’, of which some are shown in the graph. Going clockwise, firstly, two ⊖29 polymerases (at 10 units per 20 μL reaction) were compared using 5 mM random hexamer primers, including three phosphorothioate (PT) bonds on the ‘3 end, while following the suppliers’ recommendations regarding duration and temperature of the amplification step (orange section). Next, different primers (at 5 mM concentration) were assessed for their suitability (red section). The specific DNA primer (more information in the method section, as well as vector maps in the supplementary information) was designed to one site of the selected plasmid. Furthermore, we tested whether the presence of phosphatase would benefit the amplification (blue section). In four conditions, a range of three orders of magnitude were covered (0 units, 0.01 unit, 0.1 unit, and 1 unit). (**b**) Comparing the output RCA product mass (left y-axis, highlighted in green bars) to the input mass of plasmid DNA (plotted on the x-axis). Amplification factors (right y-axis, depicted with black/white dots connected by black line) versus input mass of plasmid DNA is also shown. Error bars reflect standard deviation over replicates of three for each condition. (**c**) Visualisation of nickase-treated RCA product (lane 7 and 8 (left to right)) in comparison to the presumably supercoiled input plasmids (‘PF-Nbb102-CAM’ and ‘DF-Nbb102-CAM’ in lanes 2 and 3, respectively) and the non-nickase treated linear RCA-products (lanes 5 and 6). The RCA reaction was performed using random DNA PT- and RNA hexamers (lanes 5 and 7 (PT-DNA) and 6 and 8, respectively). The dashed line indicates that the image has been cropped for simplicity. The original version can be found in Figure S3. (**d**) Beyond establishing the parameters for DNA amplification and transformation into *E. coli*, we sought a better understanding of the transformation efficiencies (electroporation and chemical) over the *in vitro* process of library assembly. The plasmid ‘PF-Nbb102-CAM’ was purified from an overnight culture and transformed either directly (condition 1 and 4), or amplified by PCR for subsequent re-insertion of the VHH encoding sequence by golden-gate assembly. The purified golden-gate product was then either transformed directly (condition 2 and 4) or subjected to RCA amplification and nicked (using Nb.BbvCI) and subsequently transformed (condition 3 and 6).

Upon confirmation of our workflow in the context of purified plasmid DNA, we next investigated the best conditions for the amplification of DNA in a ‘library context’, *i*.*e*., amplification of a golden-gate assembly. Amplification of assembled DNA was readily achieved with >18,000-fold amplification yield, generating 9.2 μg of DNA from 0.5 ng of input DNA, corresponding to one golden-gate reaction containing 8.8 x 10^8^ DNA molecules (Table 1). Post nickase treatment and purification a yield of ∼3 μg was achieved, representing an input to transformable output amplification yield of ∼6,500 fold. In comparison, a similar PCR-based amplification workflow achieved an amplification yield ∼10 fold lower than the RCA workflow. The RCA workflow is readily scalable in terms of initial volume and output mass, *e*.*g*., parallelised amplification of 40.2 ng golden-gate assembled product resulted in >700 μg purified, nickase-treated plasmid target DNA. We noticed that treatment of the golden-gate product with exonuclease to remove unassembled insert DNA was not necessary (Figure S1).

**Table 1.**
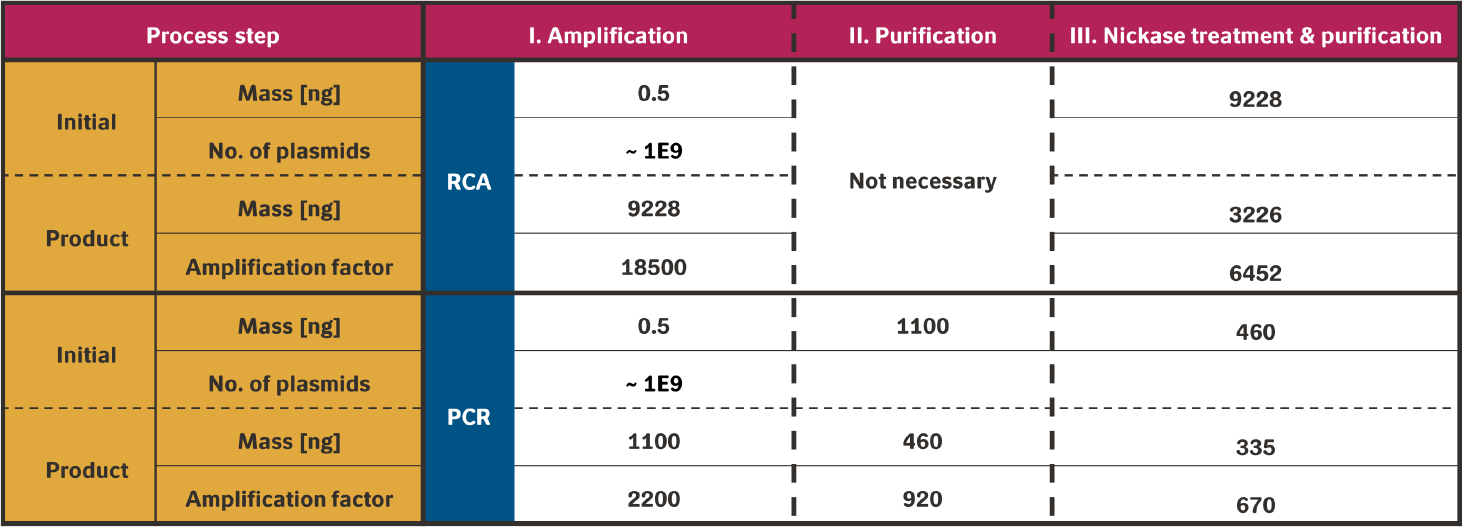
Quantification of DNA library amplification. Overview and comparison of the RCA and PCR amplification procedures by mass, total number of plasmids, and amplification factor. The values of mass are based on mass-over-volume concentration measurements, while the number of total plasmids, and amplification factor are derived from the mass values and are, therefore, dependent on each other. Importantly, the number of theoretical plasmids in the RCA input does not reflect any measurement, but rather provides a quantification of the order of magnitude. The raw measurements for mass-over-volume concentrations and subsequent calculations are provided in the supplementary information. Moreover, the calculation of the molecular mass was executed using the web application NEBioCalculator (v1.15.4 May 23, 2023) and might not reflect natural isotopic abundances. For simplicity, replicates and standard deviations are not displayed. However, the respective information can be found in the supplementary information.

**Table 2.** Highlighting the step-wise procedure of chemical transformation of the tested methods: TSS-HI, Inoue, and Zymo transformation kit.

| Step | TSS-HI protocol | Inoue protocol | Zymo protocol (- heat shock) |
| --- | --- | --- | --- |
| Preparation of DNA suspension | Pre-mix KCM buffer* (5 μL) to DNA and dH<sub>2</sub>O to a total of 25 μL | - | |
| Addition of DNA to cell suspension | Mix gently with 25 μL of competent cells | Add 1 μL of DNA | Add 1 μL of DNA |
| Incubation | Incubate for 30 min on ice | Incubate 30 min on ice | Incubate 30 min on ice |
| Heat shock | - | Heat shock for 45 sec @ 42 °C | - |
| Incubation | - | Incubation on ice (2 min.) | - |
| Addition of recovery media | Add 250 μL fresh LB | Add 1000 μL SOB + 2% glu + 5 mM MgCl<sub>2</sub> | Add 1000 μL SOB + 2% glu + 5 mM MgCl<sub>2</sub> (pre-warmed to 37 °C) |
| Recovery | Incubate for 60 min at 37 °C while shaking | Incubate for 60 min at 37 °C while shaking | Incubate for 60 min at 37 °C while shaking |
| Plating | Ready for plating in 1/10 dilution series | Ready for plating in 1/10 dilution series | Ready for plating in 1/10 dilution series |

To establish the applicability of our approach for creating large libraries, we then looked at the transformation efficiencies of circular supercoiled plasmids, relaxed circular golden-gate assembled plasmids, and linear nickase-treated RCA product across both electroporation and chemical transformation of XL-1 Blue *E. coli* cells. Here, we observed that the chemically competent cells were transformed more efficiently with our linaer nicked DNA than relaxed circular DNA (golden-gate product), yet the opposite was true for electrocompetent cells (Figure 2D). Due to this observation, we explored a variety of parameters around chemical transformation, including cell type, transformation buffers, cell concentrations, storage media, growth media for competent cell production, and growth media for post-transformation selection plates, as well as the potential scalability of chemical transformations through parallelisation, and the use of increased cell volumes (Figure S2). The most efficient in our hands was the use of a modified Inoue protocol, XL-1 blue cells grown in SOB media, stored in DMSO and plated on SOC agar supplemented with 2% Glucose and 5 mM MgCl_2_.

### RCA-mediated amplification of plasmid DNA for *S. cerevisiae* transformation

After establishing the use of the RCA nickase workflow for *E. coli*, we also tested its applicability to other common organisms used for directed evolution. We chose to investigate *S. cerevisiae*, notably the EBY100 strain for two reasons; firstly, EBY100 is commonly used for yeast surface display, a powerful technique for the ultra-high throughput quantitative screening of binder libraries^20,21^, and secondly, because yeast is known to possess strong homologous recombination activity enabling transformation with linear DNA molecules, we expected that our homology-based cloning approach may have an effect on transformation efficiencies.

*S. cerevisiae* cells are often transformed with linear DNA possessing double-stranded homologous ends; thus, we created a new homology cassette in the yeast plasmids alongside our previously described nickase-single strand cassette. This new cassette contained an FspI site in the centre of a 30 or 90 base repeat sequence, such that upon FspI treatment the RCA product would resolve to monomeric units with double-stranded homologous ends (Figure 3A). We observed that the linear FspI treated double-stranded homology DNA was most efficient in chemical transformation of yeast cells, outperforming both circular supercoiled DNA and the linear nickase treated single-stranded homology DNA by at least 10-fold (Figure 3B). This result established the utility of our approach for creating large libraries in yeast, as well as providing data confirming that linear DNA transforms yeast at higher efficiency than circular DNA. As we had observed contradictory results in *E. coli* between electroporation and chemical transformation, we also tested electrocompetent yeast and observed that linearisation significantly improved transformed efficiency versus circular DNA (Figure 3C).

**Figure 3.**
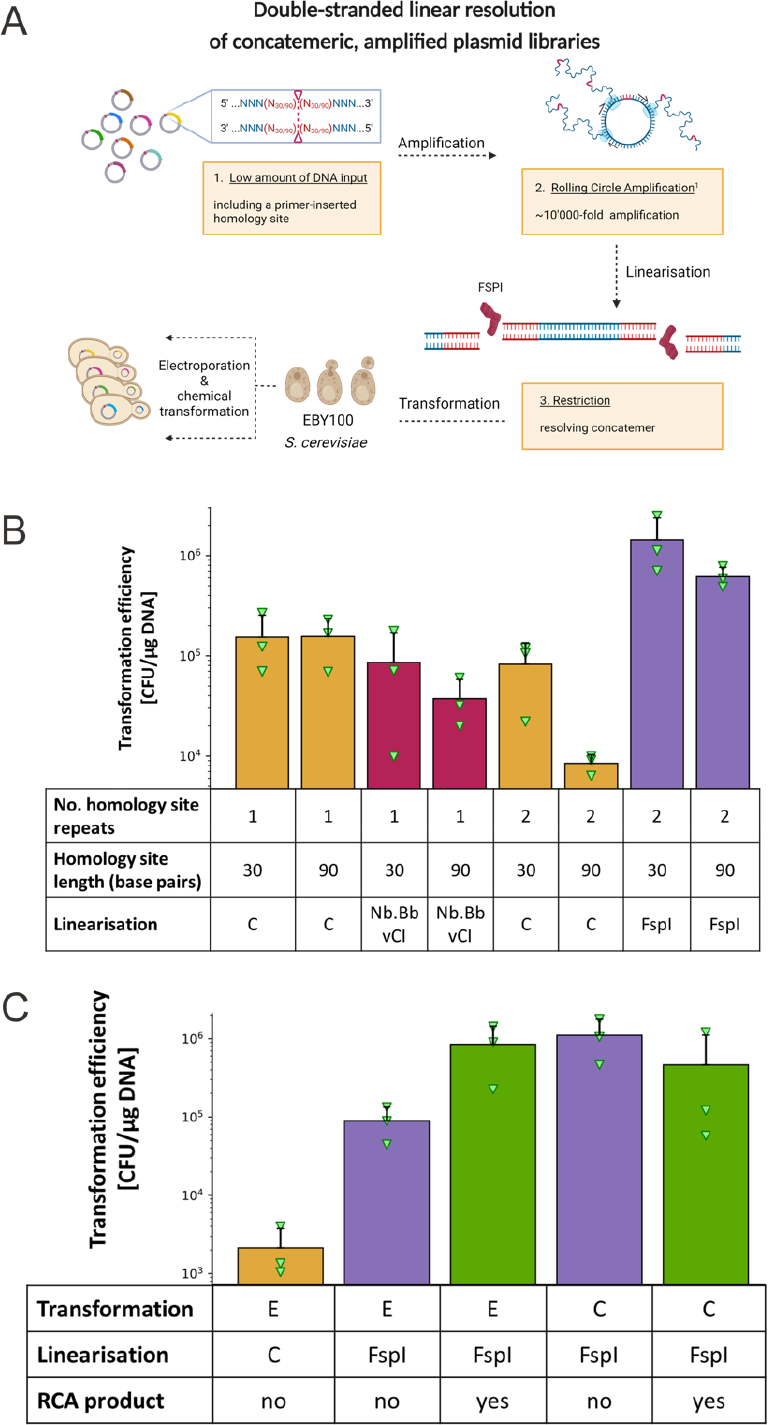
Testing DNA plasmid formats and modifications for optimal transformation into *S. cerevisiae* EBY100. **(a)** Schematic of optimised post-assembly library amplification for transformation into *S. cerevisiae*. (**b**) Using chemical transformation: Transformation efficiencies in *S. cerevisiae* across different formats (circular (orange), linear single-stranded (red), and linear double-stranded (purple)) and two different homology site lengths (30 and 90 base pairs) of similar plasmids (‘pCT-antiGFP’). Like transformations in *E. coli* strains, we tested the utility of linearised DNA for chemical transformation in *S. cerevisiae* strain EBY100. While the treatment of Nb.BbvCI resulted in a linear format with single-stranded 3’ overhangs, the double-stranded homology sites restricted by treatment with FspI leaves blunt ends. Vitally for the double-stranded homology, the according plasmid contains a repeat of the homology site which enables *in vivo* processing and assembly. (**c**) Transformation of RCA product (green) in chemical and electroporation transformation using 2 μg and 8 μg per sample, respectively, of linearised plasmid with double-stranded 90 bp homology sites.

## Discussion

The creation of large libraries is a difficult, resource and time-consuming process that significantly limits the accessibility of directed evolution to the wider scientific community. Whilst novel technologies, such as *in vivo* hypermutation (*e*.*g*., in *E. coli* as PACE^22^/PRANCE^23^ or MutaT7^24^, *B. thuringiensis* as BacORep^25^ and *S. cerevisiae* as OrthoRep^26^), can circumvent the need to physically transform as many cells as you want your library size to comprise, it is still likely that physical transformation will remain the method of choice for many researchers due to specific library designs, or the choice of screening technologies which are not compatible with the mentioned technologies. In addition, an increased transformant yield is also useful in other fields, where subsequent *in vivo* mutation is undesirable, such as basic biology, sequencing, DNA storage, etc. As such, any methods that can simplify the creation of large libraries are of great use. Interestingly, it has been suggested that transformation of *E. coli* is not as clonal as typically considered, thus in the creation of large libraries it is likely that the diversity screened is actually greater than the number of transformants^27^. Such effects can be resolved after candidate identification through plasmid extraction and retransformation at lower DNA:cell ratios, and in the case of phage display would be resolved after the output of the first round of panning is used to infect cells (presuming a multiplicity of infection <1).

In this work, we have developed a simple protocol for the creation of large libraries in *E. coli* through chemical transformation. Whilst it was an unintended consequence of the interesting molecular biology that we uncovered, (that linear DNA does not transform efficiently via electroporation), the outcome of focussing on chemical transformation is elegant as it is the least limited by accessibility to specific pieces of equipment. The observed difference in efficiency between linear DNA transformation into electrocompetent cells and chemically competent cells can likely be explained by the unique mechanisms by which DNA passes into the cells of each approach. However, as the exact mechanisms of either chemical competency or electrocompetency are not fully understood, it is difficult to posit much beyond this.

To further extend the utility of our approach, we confirmed its use for large library creation in *S. cerevisiae*, a highly useful organism in biotechnology and synthetic biology for which RCA-mediated library amplification has not been explored previously. We expect that the product of our protocol could be of benefit in organisms other than *E. coli* or *S. cerevisiae*, such as the rapidly developing biotechnology chassis organism *V. natriegens*^28^ for which efficient and scalable natural transformation techniques have been established^29^. Additionally, the linear DNA product of our protocol could be of use for genome integration rather than plasmid formation.

In conclusion, our workflow can readily yield ∼6,500-fold amplification of purified transformable DNA, such that large libraries (>10^8^) can easily be achieved from even single small-scale (nanogram) library assembly reactions in both *E. coli* and *S. cerevisiae*. This is achieved in as simple a manner as possible, with a focus on reduction of both cost and resource usage.

## Materials and methods

### Plasmids and primers

Plasmids employed for transformations in *E. coli* are derivatives of the plasmid ‘PF-Nbb102-CAM’, a VHH encoding plasmid (phagemid-derived from the commercially available pADL22c) with chloramphenicol resistance gene, and have been modified by amplification of designed primers ordered from IDT (Integrated DNA Technologies, Inc.). Similarly, *S. cerevisiae* transformation tested were conducted by employing derivates of the plasmid ‘pCT anti-GFP’ a yeast display vector containing an anti-GFP VHH

Sequences are provided in Supplementary table 1.

### DNA manipulation

#### 1. Nickase site insertion

In the phagemid context nickase sites were added through PCR amplification using Q5 DNA polymerase and appropriate DNA primer pairs (F NbBbvCI 10bp ‘gcaggttt*gctgagg*CCCGACTGGAAAGCGGGC’ with R NbBvCI 10 bp ‘gcaaacct*gctgagg*GTCGTGCCAGGGCATCCC’ and F NbBbvCI 15 bp ‘gcacgacaggttt*gctgagg*CCCGACTGGAAAGCGGGC’ with R NbBbvCI 15 bp ‘gcaaacctgtcgt*gctgagg*CCAGGGCATCCCTCCTTTCA’). In upper case is the annealing site, italics the Nb.BbvCI site, and underlined the homology region for *in vivo* assembly. PCR product was DpnI treated and purified using homemade SPRI beads (1 mL Sera-Mag™ Carboxylate-Modified [E3] Magnetic Particles: 44152105050250 in 50 mL 2.5 M NaCl, 20% PEG-8000) in a 1:1 volumetric ratio followed by magnetic capture and washing with 70% Ethanol, Tween-20 0.05%, removal of excess ethanol and elution in MilliQ water. Transformation of these linear DNA products results in plasmids containing the appropriate nickase cassettes through *in vivo* assembly.

Similarly the yeast plasmid nickase cassettes were inserted through amplification of the pCT anti-GFP backbone with appropriate DNA primer pairs followed by transformation into *E. coli* for *in vivo* assembly (F Nick 30 bp ‘tggccgattcattaatgcagttt*gctgagg*CTCCAATTCGCCCTATAGTG’ with R Nick 30 bp ‘ctgcattaatgaatcggccacct*gctgagg*CTCAATTCTCTTAGGATTCGATTC’ and F Nick 90 bp ‘tacactattctattggaatcttaatcattctggccgattcattaatgcagttt*gctgagg*CTCCAATTCGCCCTATAGTG’ with R Nick 90 bp ‘gattccaatagaatagtgtataaattatttatcttgaaaggagggatgcccct*gctgagg*CTCAATTCTCTTAGGATTCGATTC’). In upper case is the annealing site, italics the Nb.BbvCI site, and underlined the homology region for *in vivo* assembly. The double-stranded homology cassettes were built on top of the pCT anti-GFP plasmids already containing the 30 bp or 90 bp nickase cassettes, firstly the appropriate plasmids were amplified with (F pCT BsaI ‘gagtag*ggtctc*cTGAGGCTCCAATTCGCC’ and R pCT BsaI ‘gaggat*ggtctc*aCTCAGCAAACTGCATTAATGAATCGGCCA’), secondly the 30 bp/90bp cassettes were created by mixing the two appropriate primers alone in a PCR reaction (F FspI cassette 30 bp ‘gaggta*ggtctc*attgcgcaggTGGCCGATTCATTAATGCAG’ with R FspI cassette 30 bp ‘gaggat*ggtctc*actcagcaaaCTGCATTAATGAATCGGCCA’ and F FspI cassette 90 bp ‘gagat*ggtctc*attgcgcaggggcatccctcctttcaagataaataatttaTACACTATTCTATTGGAATC’ with R FspI cassette 90 bp ‘gagat*ggtctc*actcagcaaactgcattaatgaatcggccagaatgattaaGATTCCAATAGAATAGTGTA’. In upper case is the annealing site, italics the BsaI site. The products were then appropriately mixed in a golden-gate assembly reaction using BsaI-HFv2 (NEB: R3733L) and T4 DNA ligase (NEB: M0202L) and transformed into *E. coli*.

#### 2. Amplification

Initially, while seeking to optimise the conditions for amplification, we tested two different phi29 polymerases: NEB phi29 polymerase (New England Biolabs, catalogue no.: M0269L) and EquiPhi29 DNA Polymerase (ThermoFisher Scientific, catalogue no.: A39390). Employing both polymerases, we assessed three randomised hexamer primers (random DNA hexamers (NNNNNN), random DNA hexamers with 3x phosphorothioate bonds (N*N*N*NNN), and random RNA hexamers) and one defined primer pair, which was known to anneal to specific sites of the plasmid in question (forward-PD116 ‘CATGACCAAAATCCCTTAAC’ & reverse-PD117 ‘CATGAGCGGATACATATTTG’).

In general, <1 ng of DNA of interest was pipetted to a mix of 1x rCutsmart buffer (New England Biolabs), 100 μM of primer, and nuclease-free water. The sample denatured at 95 °C for 3 min. and immediately placed on ice for another 3-5 min. For the amplification step, 1 mM dNTP, 1x rCutsmart buffer, MilliQ, and 10 U of the respective ⊖29 DNA polymerase were added to a final reaction volume of 20 μL.

For amplifications facilitated by EquiPhi29 DNA polymerase, we added 1 mM DTT and incubated the reaction mixed at 45 °C for 3 hours. Reaction mixes containing NEB Phi29 0.1 mg/mL recombinant albumin were incubated for 16 hours at 30 °C. Both polymerases were heat inactivated by heating the sample at 65 C for ten minutes.

#### 3. Linearisation

To resolve the concatemeric structure, 1 μg RCA product was restricted by employing 10 U nicking endonuclease Nb.BbvCI (New England Biolabs, R0631L) in a 50 μL reaction volume with 1x rCutsmart buffer for 1 h at 37 °C, followed by a heat inactivation step at 80 °C for 20 min. In case of double-stranded linear DNA applied in transformation experiments with *S. cerevisiae*, the RCA product was resolved to linear monomers by restriction using the endonuclease Fsp1 (New England Biolabs, R0135L) under the similar conditions.

All DNA concentration measurements were conducted with the Qubit broad range assay kit (ThermoFisher Scientific Inc.)

### Production of competent cells

#### 1. Preparation of electrocompetent *E*. *coli* strains XL-1 Blue and BL21 (DE3)

Five mL LB medium (including 15 mg/mL tetracycline for XL-1 Blue) were inoculated from respective glycerol stocks and grown overnight at 37 C and 200-250 rpm. Next, 500 mL of SOB (pH 7.5, 5 g/L yeast extract, 20 g/L tryptone/peptone, 0.584 g/L NaCl, 0.186 g/L KCl, 2.4 g/L MgSO4) were added to a 2.5 L shake flask and inoculated by cells from the previous culture to an initial OD of 0.007. The culture grew slowly at 20 °C and 250 rpm for 42 h to a final OD of 0.6.

Subsequently, the culture was stored on ice, pelleted (2500 g, 4 °C, 9/12/15 minutes), and washed with ice-cold MilliQ water three times, while steadily reducing the resuspension volume (250/50/25 mL) and adding 10% final concentration of glycerol from the second wash step on-wards. Lastly, pellets were resuspended in 5 mL ice-cold MilliQ water with 20% glycerol in case the cells were stored in -80 °C or without glycerol if directly used for transformation.

#### 2. Preparation of chemical competent *E*. *coli* strains XL-1 Blue, and BW25113

a. Adjusted Inoue protocol Cultures from *E. coli* cell strain XL-1 Blue (no antibiotic resistance gene) were grown as previously described for the preparation of electrocompetent *E. coli* and harvested at an OD between 0.4 and 0.6. The cultures were pelleted (4000 g, 10 minutes, and 4 °C) and resuspended in sterilised 250 mL TB buffer (pH 6.7 (adjusted with KOH), PIPES 3.021 g/L, CaCl_2_*2H_2_O 11.025 g/L, KCl 18.637 g/L, MnCl_2_.4H2O 10.885 g/L). Pelleting was repeated after storing the resuspended mixture on ice for 10 minutes. Next, cells were resuspended in 40 mL TB buffer and 3 mL DMSO, gently mixed, and stored on ice for another 10 minutes. Subsequently, the resuspended cells were distributed in aliquots of 100 μL and either directly used or stored at -80 °C.
b. TSS-HI (transformation storage solution optimized by Hannahan and Inoue method) To test further protocols for chemical competence for *E. coli*, we took inspiration from Yang et al. and tested the strain BW25113^30^ (no antibiotic resistance) with their improved TS-HI protocol^31^. 4 mL of LB medium were inoculated and cells grown overnight at 250 rpm and 37 C. 1 % (40 μL) of the culture were transferred to fresh 50 mL LB and grown to an OD (600 nm) of 0.5. Subsequently, the cells were stored on ice for 10 min., centrifuged at 4 °C and 4000 g, and resuspended 1 mL chilled (0 °C) TSS-HI.
c. Mix & Go *E. coli* Transformation Kit (T3001) Benchmarking the chemical competence protocols for the *E. coli* strain XL-1 Blue, we tested the commercial transformation kit ‘Mix & Go *E. coli* Transformation Kit (T3001)’ available at (zymoresearch.com) and followed the general guidance (V.1.18) including ‘Notes for High Efficiency Transformation’.

#### 3. Chemical competent *S*. *cerevisiae* strain EBY100

Testing the utility the robust workflow of DNA amplification and nickase treated linear homology sites could provide for yeast transformation, we prepared chemical competent *S. cerevisiae* strain EBY100 cells following the Li/Ac protocol of Gietz & Schiestl^4^. Deviating from the protocol we inoculated 100 mL of the second culture with a final OD_600_ of 0.25. Cells were harvested after 6 h of incubation at an OD_600_ of 0.95, representing a total cell titer of 9.5 * 10^8 cells in 100 mL culture volume.

#### 4. Electroporation competent *S*. *cerevisiae* strain EBY100

For the preparation of electroporation competent EBY100 cells, we followed the protocol of Benatuil et al.^32^. In short, we grew *S. cerevisiae* cells (EBY100) overnight to stationary phase (OD_600_ ∼3) in YPD media (10 g/L yeast nitrogen base, 20 g/L Peptone and 20 g/L D-(+)-Glucose) on a platform shaker at 250 rpm and 30 °C. The next morning, 500 mL fresh YPD media was inoculated using an aliquot of the overnight culture to an initial OD_600_ of 0.3 and subsequently incubated at 30 °C and 225 rpm until OD_600_ was 1.6. Cells were collected by centrifugation at 3000 rpm for 3 min. at 4 °C, washed twice with 250 mL ice-cold water and once 250 mL of ice-cold electroporation buffer (1 M Sorbitol / 1 mM CaCl2). Next, follows conditioning of cells by re-suspending the cell pellet in 100 mL 0.1 M LiAc/10 mM DTT and shaking at 230 rpm in a culture flask for 30 minutes at 30 °C. Once again, cells were collected, washed once by 250 mL ice-cold electroporation buffer, and re-suspended the cell pellet in 500 to 1000 μL electroporation buffer to reach a final volume of 4.0 mL. This corresponds to approximately 8 × 10^9 cells in total and is sufficient for 10 electroporations at 400 μL total volume per 0.2-m gap cuvette. The cells were kept on ice until electroporation.

### Transformation

#### 1. Electroporation for *E*. *coli* strains

Aliquots of electrocompetent *E. coli*, directly after preparation or stored at -80 °C, were slowly thawed on ice. Once thawed, the sample DNA (in MilliQ) was added to a maximal voluminal ratio of 5%, typically 5 μL or less to 100 μL of cell suspension, gently mixed by pipetting, and incubated for 3-5 min on ice. Next, the chilled DNA/cell suspension was filled into a pre-chilled 0.1-cm gap cuvette (BioRad) and electroporated using a Gene Pulse Controller electroporation system (Bio-Rad) at 1.5 kV voltage, 200 Ω resistance, and 25 kF/cm^2^ capacity. After applying the electroshock (usually response time around 4-5 ms), we rescued the cells by pipetting 1 mL of pre-warmed (37 °C) SOC media including 2% Glucose and 5 mM MgCl_2_ into the cuvette, slowly up- and down, and then transferring to a 1.5 mL reaction tube. The suspension was incubated for 90 minutes, at 37 °C, 750 rpm on a table-top shaker. In the meantime, selective plates (SOC, 2% glucose, and respective antibiotics) were pre-warmed to 37 °C. Lastly, the samples were diluted in a 1/10 dilution series and plated as 5 μL spots in triplicates on SOC agar plates including 2% glucose and chloramphenicol selectivity.

#### 2. Chemical transformation for *E*. *coli* strains

Generally, the chemical transformation for all three tested methods is rather similar. In order to depict similarities and differences the transformation procedures can be found table x.

#### 3. Chemical transformation for *S*. *cerevisiae* strain EBY100

Yeast transformations were always conducted directly subsequent to cell preparation. Aliquots of 100 μL cell suspension were spun down at 13000 x rpm for 30 s and resuspended in 336 μL transformation mix (0.1 M LiAc, 240 μL PEG 3350 50% (w/v)) including 0.28 mg/mL single-stranded carrier DNA (Dual Systems, lot n. 6001120319) and the specific DNA sample of interest. Samples were exposed to a subsequent 40 min. heat shock at 42 °C and 650 rpm shaking. Followed by pelleting and resuspending in 1 mL sterile MilliQ water, samples were subjected to a 1/10 dilution series and plated on selective tryptophan-dropout media agar plates.

#### 4. Electroporation for *S*. *cerevisiae* strain EBY100

Electroporation of yeast cells followed their preparation immediately. The cells suspension is aliquoted to 370 μL per sample and mixed gently with 30 μL MilliQ including 8 μg target DNA. After an incubation not exceeding five minutes the chilled DNA/cell suspension was filled into a pre-chilled 0.2-cm gap cuvette (BioRad) and electroporated using a Gene Pulse Controller electroporation system (Bio-Rad) at 2.5 kV voltage, 200 Ω resistance, and 25 kF/cm^2^ capacity. After applying the electroshock (usually response time around 3.8-4.5 ms), we rescued the cells by pipetting 1 mL of pre-warmed (30 °C) 1:1 mix of 1 M sorbitol : YPD media into the cuvette, slowly up- and down, and then transferring to a 50 mL centrifuge tube, which already contained 6 mL pre-warmed 1:1 mix of 1 M sorbitol:YPD media. The rescue procedure was repeated with a fresh 1 mL, yielding a total of 8 mL culture. The suspension was incubated for 60 minutes, at 30 °C, and at 225 rpm. In the meantime, selective plates (SD (-Trp) + 2% Glucose including 10 mg/mL chloramphenicol) were pre-warmed to 30 °C. Lastly, we diluted the samples in a 1/10 dilution series and plated 30 μL of each condition in one quarter of a selective plate. Colonies were counted on the second day after plating.

## Author Contributions

- Idea & concept: <u>TF;</u>
- Supervision: <u>TF</u>, CA, TPJ;
- Funding: <u>CA, AHL</u>;
- Experimental work: <u>DSW</u>, TF, MDO;
- Data analysis, editing, and interpretation: <u>DSW, TF</u>;
- Contributed to protocol optimisation: <u>TF</u>, DSW, MDO, ES;
- Drafting manuscript including graphic representations: <u>TF, DSW</u>, TPJ;
- Internal review and submission: <u>TPJ, TF</u>, CA, MDO, ES, AHL;

All authors contributed to the manuscript and approved its final version.

## Data Availability

All underlying raw data files and calculations are accessible.

