## Supplementary information for "Simplified post-assembly plasmid library amplification for increased transformation yields in *E. coli* and *S. cerevisiae*"

**SUPPLEMENARY INFORMATION**

Supplementary figures and tables:

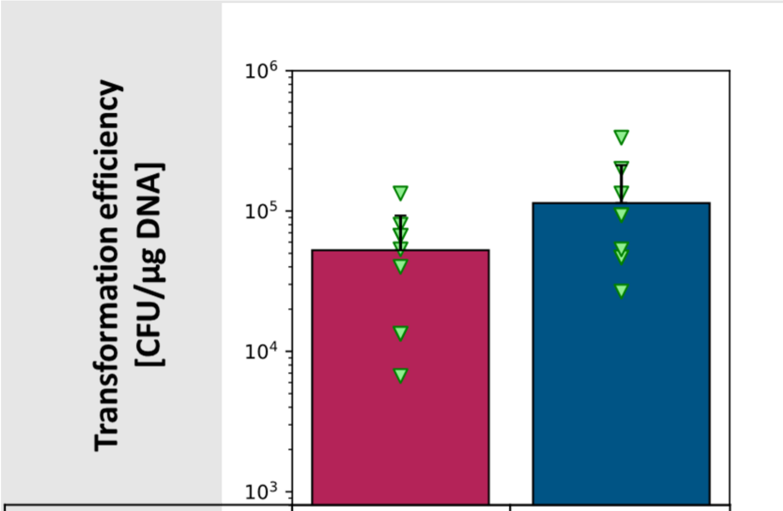


| | | |
|---|---|---|
| **Exonuclease** | yes | no |
| **Linearisation** | yes | yes |
| **RCA product** | yes | yes |

**Supplementary figure 1. Effect of exonuclease treatment in the process of library amplification using RCA on subsequent transformation efficiency.** To further optimise the library creation process, we assessed whether treatment with exonuclease V (NEB, Catalogue-no.: M0345L), a RecBCD complex from *E. coli*, would have an effect on transformation efficiency. Exonuclease V bidirectionally hydrolyses phosphodiester bonds between nucleotides in linear double-stranded DNA, leaving ideally only assembled DNA after golden-gate assembly of the VHH-containing insert into a suitable vector ('Pf-Nb-b102-Cam-lin10'). From the calculated transformation efficiency of the individual samples and their replicates (depicted in green triangles), average (highlighted by red (including exonuclease treatment) and blue bars), and standard deviation were noted. Following the manufacturer's instructions, 10 units of exonuclease V were employed per 1 µg of DNA, while adding 1 mM ATP. Reaction was quenched after 30 min by adding >10 mM EDTA. Error bars represent standard deviation of replicates nand face only upwards for simplicity. Individual replicates are highlighted in green triangles.

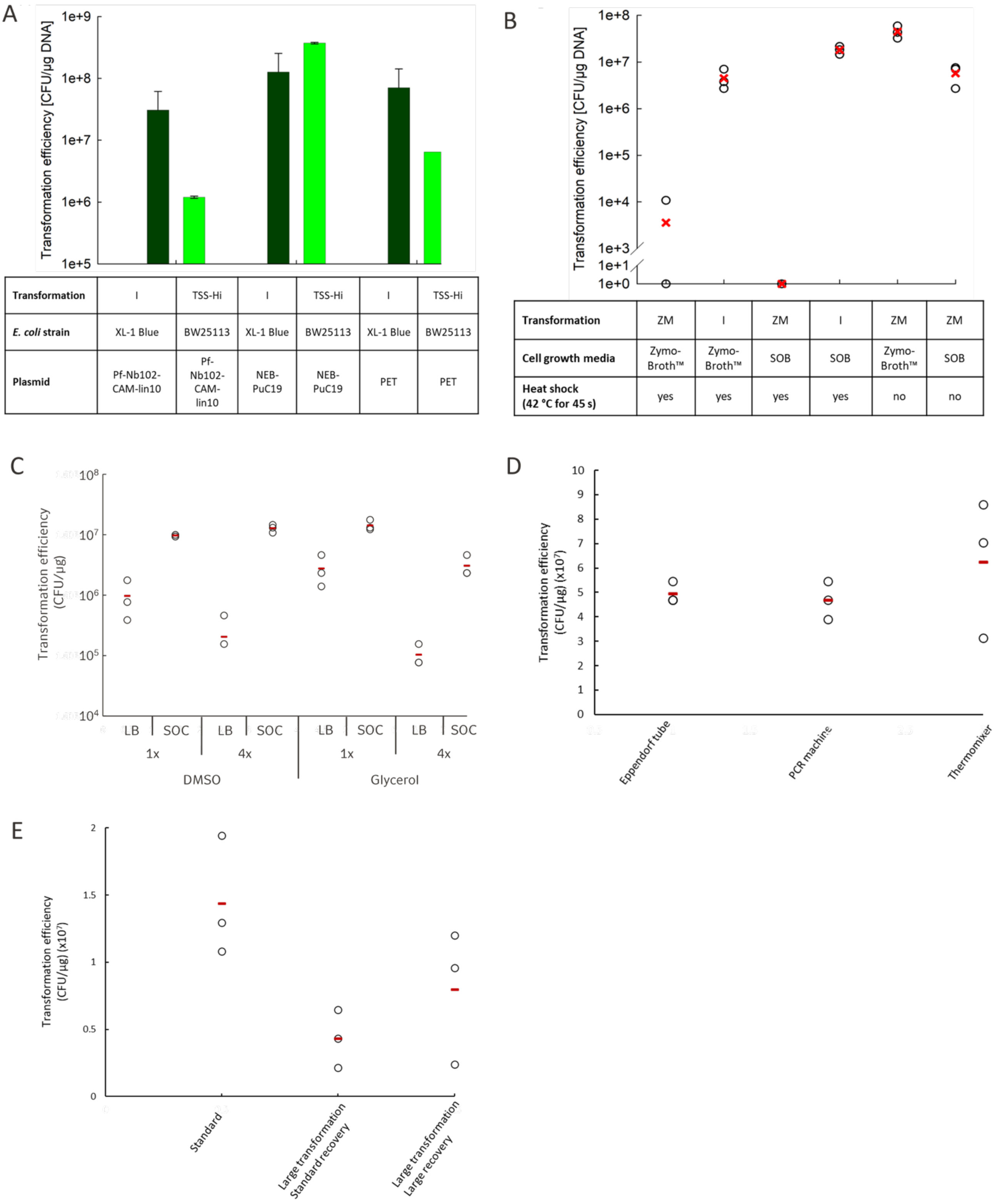


| Transformation | I | TSS-Hi | I | TSS-Hi | I | TSS-Hi |
|---|---|---|---|---|---|---|
| *E. coli* strain | XL-1 Blue | BW25113 | XL-1 Blue | BW25113 | XL-1 Blue | BW25113 |
| Plasmid | Pf-Nb102-CAM-lin10 | Pf-Nb102-CAM-lin10 | NEB-PuC19 | NEB-PuC19 | PET | PET |

| Transformation | ZM | I | ZM | I | ZM | ZM |
|---|---|---|---|---|---|---|
| Cell growth media | Zymo-Broth™ | Zymo-Broth™ | SOB | SOB | Zymo-Broth™ | SOB |
| Heat shock (42 °C for 45 s) | yes | yes | yes | yes | no | no |

**Supplementary figure 2. Benchmarking chemical transformation methods. (a)** Seeking the best performance, the current method of chemical *E. coli* transformation, named 'I' referring to 'Inoue' in dark green, was evaluated in transformation efficiency over the range of three different plasmids against the 'TSS-HI' method depicted in light green. (**b**) Next, 'I' was tested against the commercial *E. coli* transformation kit Zymo Mix&Go (Zymo Research, catalogue-no.: T3001). Most noticeably, a heat shock procedure (heating to 42 °C for 45 sec. and immediate cooling on ice) is not included in the protocol of Zymo research. We, therefore, tried different combinations of transformation protocol, media, and heat shock. (**c**) Chemically competent XL-1 Blue cells were prepared as per the Inoue protocol, with differences in their final storage formulation (DMSO or glycerol), how concentrated the aliquots were (with 4x denoting a resuspension in a 4-fold lower final volume), and what agar the transformations were plated on (LB Cam, or SOC 2% Glucose Cam). (**d**) Parallelisation of transformations was explored by comparing standard protocols (in an eppendorf tube) to use of a 96-well PCR plate containing aliquots of cells and heat-shock in a PCR machine (programmed to 4 °C for 30 minutes, 42 °C for 45 seconds, and 4 °C for 5 minutes) or in 96-well metallic cold blocks on ice and a 96-well thermomixer programmed to 42 °C (**e**) The volumetric scale-up of transformation was explored by comparing standard transformations in Eppendorf tubes (100 µL cells + 900 µL recovery media) to transformation of 1 mL cells in an Eppendorf tube (large transformation) and either standard recovery (100 µL cells + 900 µL recovery media) or large recovery (900 µL cells + 9000 µL recovery media). Data are the mean of triplicates (red) with individual data points indicated.

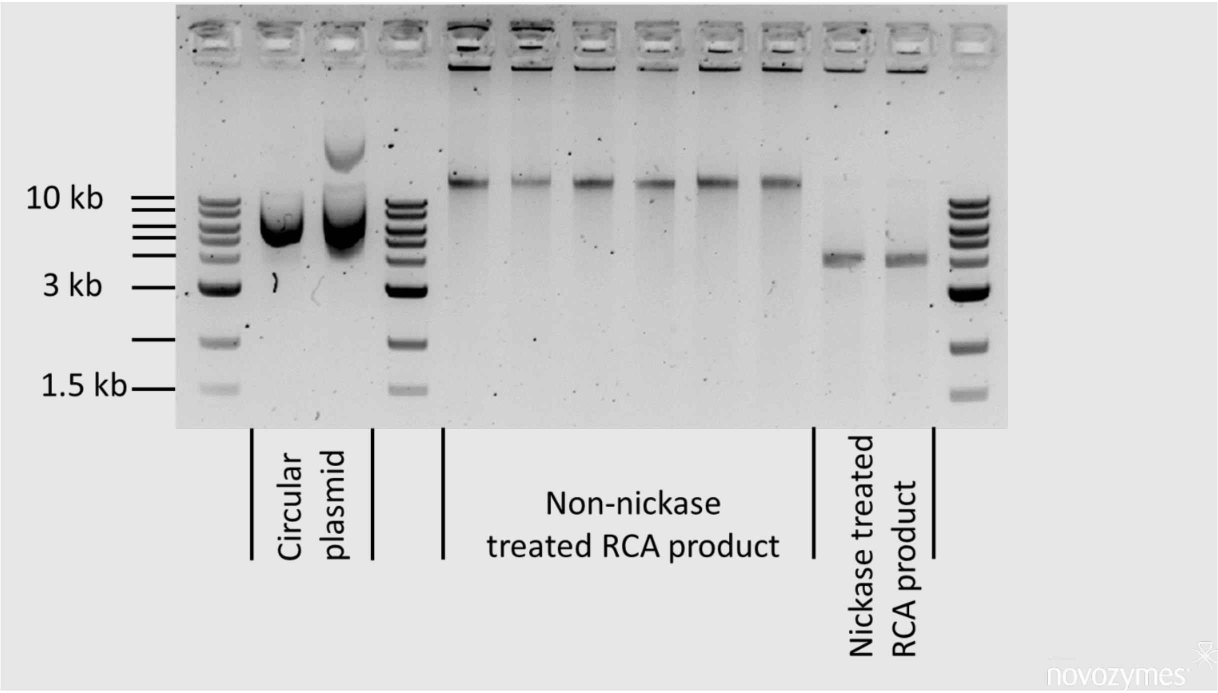


**Supplementary figure 3. Nickase-mediated resolution of concatermeric, RCA amplified plasmid.** Please note that this is the same image as in Figure 2C, but not cropped. Visualisation of nickase-treated RCA product (lane 11 and 12 (left to right)) in comparison to the presumably supercoiled input plasmid ('PF-Nbb102-CAM' and 'DF-Nbb102-CAM' in lane 2, respective 3) and the non-nickase treated linear RCA-products (lanes 5-10). The RCA reaction was performed using random DNA PT- and RNA hexamers (lane 5, 7, 9, and 11, and 6, 8, 10, and 12, respectively). Furthermore, to obtain an accurate comparison to the nickase-treated samples, the RCA product in lane 9 and 10 underwent the same treatment of buffers, heat, and additional purification without addition of the nickase enzyme Nb.BbvCI as the nickase treated samples in lane 11 and 12. Samples in lane 5 and 6 were not purified after the RCA reaction, while the samples in lane 7 to 10 were purified using SPRI beads.

**Supplementary table 1: Plasmid sequences**

| Plasmid | Sequence 5'-3' (relevant cassette region in bold and uppercase, with restriction site(s) underlined) |
|---|---|
| pCT-antiGFP Nick 30 | ccaatacgcaaaccgcctctccccgcgcgttggccgattcattaatgcagctggcacgacaggtttcccgactggaaagcgggcagtgagcgcaacgcaatt<br>aatgtgagttacctcactcattaggcaccccaggctttacactttatgcttccggctcctatgttgtgtggaattgtgagcggataacaatttcacacaggaaacag<br>ctatgaccatgattacgccaagctgccagatctgcagccgctatatatccgcggattaacgcgctagccggctgggccgtcgaacggaattaaccctcactaa<br>agggaacaaaagctgggtacccgacaggttatcagcaacaacacagtcatatccattctcaattagctctaccacagtgtgtgaaccaatgtatccagcacca<br>cctgtaaccaaaacaattttagaagtactttcactttgtaactgagctgtcatttatattgaattttcaaaaattcttactttttttttggatggacgcaaagaagtttaata<br>atcatattacatggcattaccaccatatacatatccatatacatatccatatctaatcttacttatatgttgtggaaatgtaaagagccccattatcttagcctaaaaaa<br>accttctctttggaactttcagtaatacgcttaactgctcattgctatattgaagtacggattagaagccgccgagcgggtgacagccctccgaaggaagactctc<br>ctccgtgcgtcctcgtcttcaccggtcgcgttcctgaaacgcagatgtgcctcgcgccgcactgctccgaacaataaagattctacaatactagcttttatggttatg<br>aagaggaaaaattggcagtaacctggccccacaaaccttcaaatgaacgaatcaaattaacaaccataggatgataatgcgattagtttttttagccttatttctg<br>gggtaattaatcagcgaagcgatgatttttgatctattaacagatatataaatgcaaaaactgcataaccactttaactaatactttcaacattttcggtttgtattactt<br>cttattcaaatgtaataaaagtatcaacaaaaaattgttaatatacctctatactttaacgtcaaggagaaaaaaccccggatcgaattccctacttcatacattttc<br>aattaagatgcagttacttcgctgtttttcaatattttctgttattgcttcagttttagcacaggaactgacaactatatgcgagcaaatcccctcaccaactttagaatc<br>gacgccgtactctttgtcaacgactactattttggccaacgggaaggcaatgcaaggagtttttgaatattacaaatcagtaacgtttgtcagtaattgcggttctca<br>cccctcaacaactagcaaaggcagccccataaacacacagtatgtttttaaggacaatagctcgacgattgaaggtagatacccatacgacgttccagacta<br>cgctctgcaggctagtggtggcggaggttctggtggaggcgggtctggtgggggaggatctgccatggtgcagctggtggaaagcggtggtgcactggtgca<br>gccaggtggcagcctgcgtctgagctgcgcggcgagcggctttccggtgaaccgctatagcatgcgctggtatcgtcaggcgccgggcaaagaacgcgaat<br>gggtggcgggcatgagcagcgcgggcgatcgcagcagctatgaagatagcgtgaaaggccgctttaccattagccgcgatgatgcgcgcaacaccgtgta<br>tctgcagatgaacagcctgaaaccggaagataccgcggtgtattattgcaacgtgaacgtgggctttgaatattggggccagggcacccaggtgaccgtgag<br>cagcgcggccgctagcgaaaccccaggtacgagcgaaagcgcgaccccggaacatatggtgccgaccatcgtgatggtggacgcctacaagcgttaca<br>agtaatagctcgagatctgataacaacagtgtagatgtaacaaaatcgactttgttcccactgtacttttagctcgtacaaaatacaatatacttttcatttctccgtaa<br>acaacatgttttcccatgtaatatccttttctatttttcgttccgttaccaactttacacatactttatatagctattcacttctatacactaaaaaactaagacaattttaattt<br>tgctgcctgccatatttcaatttgttataaattcctataatttatcctattagtagctaaaaaaagatgaatgtgaatcgaatcctaagagaattgag**CCTCAGC**<br>**AGGTGGCCGATTCATTAATGCAGTTTGCTGAGG**ctccaattcgccctatagtgagtcgtattacaattcactggccgtcgttttacaacgtc<br>gtgactgggaaaaccctggcgttacccaacttaatcgccttgcagcacatccccctttcgccagctggcgtaatagcgaagaggcccgcaccgatcgccctttcc<br>caacagttgggcagcctgaatggcgaatggacgcgccctgtagcggcgcattaagcgcggcgggtgtggtggttacgcgcagcgtgaccgctacacttgcc<br>agcgccctagcgcccgctcctttcgctttcttcccttcctttctcgccacgttcgccggctttccccgtcaagctctaaatcgggggctccctttagggttccgatttagt<br>gctttacggcacctcgaccccaaaaaacttgattagggtgatggttcacgtagtgggccatcgccctgatagacggtttttcgccctttgacgttggagtccacgtt<br>ctttaatagtggactcttgttccaaactggaacaacactcaaccctatctcggtctattcttttgatttataagggattttgccgatttcggcctattggttaaaaaatgag<br>ctgatttaacaaaaatttaacgcgaattttaacaaaatattaacgcttacaatttcctgatgcggtattttctccttacgcatctgtgcggtatttcacaccgcatagatc<br>ggcaagtgcacaaacaatacttaaataaatactactcagtaataacctatttcttagcatttttgacgaaatttgctattttgttagagtcttttacaccatttgtctccac<br>acctccgcttacatcaacaccaataacgccatttaatctaagcgcatcaccaacattttctggcgtcagtccaccagctaacataaaatgtaagctttcggggctc<br>tcttgccttccaacccagtcagaaatcgagttccaatccaaaagttcacctgtcccacctgcttctgaatcaaacaagggaataaacgaatgaggtttctgtgaa<br>gctgcactgagtagtatgttgcagtcttttggaaatacgagtcttttaataactggcaaaccgaggaactcttggtattcttgccacgactcatctccatgcagttgga<br>cgatatcaatgccgtaatcattgaccagagccaaaacatcctccttaggttgattacgaaacacgccaaccaagtatttcggagtgcctgaactatttttatatgct<br>tttacaagacttgaaattttccttgcaataaccgggtcaattgttctctttctattgggcacacatataatacccagcaagtcagcatcggaatctagagcacattctg<br>cggcctctgtgctctgcaagccgcaaactttcaccaatggaccagaactacctgtgaaattaataacagacatactccaagctgcctttgtgtgcttaatcacgta<br>tactcacgtgctcaatagtcaccaatgccctccctcttggccctctccttttcttttttcgaccgaattaattcttaatcggcaaaaaaagaaaagctccggatcaaga<br>ttgtacgtaaggtgacaagctatttttcaataaagaatatcttccactactgccatctggcgtcataactgcaaagtacacatatattacgatgctgtctattaaatgct<br>tcctatattatatatatagtaatgtcgtttatggtgcactctcagtacaatctgctctgatgccgcatagttaagccagccccgacacccgccaacacccgctgacg<br>cgccctgacgggcttgtctgctcccggcatccgcttacagacaagctgtgaccgtctccgggagctgcatgtgtcagaggttttcaccgtcatcaccgaaacgc<br>gcgagacgaaagggcctcgtgatacgcctatttttataggttaatgtcatgataataatggtttcttaggacggatcgcttgcctgtaacttacacgcgcctcgtatct |

| | |
|---|---|
| | tttaatgatggaataatttgggaatttactctgtgtttatttatttttatgttttgtatttggattttagaaagtaaataaagaaggtagaagagttacggaatgaagaaaa<br>aaaaataaacaaaggtttaaaaaatttcaacaaaaagcgtactttacatatatatttattagacaagaaaagcagattaaatagatatacattcgattaacgata<br>agtaaaatgtaaaatcacaggattttcgtgtgtggtcttctacacagacaagatgaaacaattcggcattaatacctgagagcaggaagagcaagataaaag<br>gtagtatttgttggcgatcccctagagtcttttacatcttcggaaaacaaaaactattttttctttaatttctttttttactttctatttttaatttatatatttatattaaaaaattta<br>aattataattatttttatagcacgtgatgaaaaggacccaggtggcacttttcggggaaatgtgcgcggaacccctatttgtttatttttctaaatacattcaaatatgta<br>tccgctcatgtcgagacgttgggtgaggttccaactttcaccataatgaaataagatcactaccgggcgtattttttgagttatcgagattttcaggagctaaggaa<br>gctaaaatggagaaaaaaatcactggatataccaccgttgatatatcccaatggcatcgtaaagaacattttgaggcatttcagtcagttgctcaatgtacctata<br>accagaccgttcagctggatattacggcctttttaaagaccgtaaagaaaaataagcacaagttttatccggcctttattcacattcttgcccgcctgatgaatgctc<br>acccggagttccgtatggcaatgaaagacggtgagctggtgatatgggatagtgttcacccttgttacaccgttttccatgagcaaactgaaacgttttcatcgctc<br>tggagtgaataccacgacgatttccggcagtttctacacatatattcgcaagatgtggcgtgttacggtgaaaacctggcctatttccctaaagggtttattgagaa<br>tatgtttttcgtctcagccaatccctgggtgagtttcaccagttttgatttaaacgtggctaatatggacaacttcttcgcccccgttttcaccatgggcaaatattatacg<br>caaggcgacaaggtgctgatgccgctggcgattcaggttcatcatgccgtttgtgatggcttccatgtcggcagaatgcttaatgaattacaacagtactgcgatg<br>agtggcagggcggggcgtaatttttttaaggcagttattggtgcccttaaacgcctggtgctacgcctgaataagtgataataagcggatgaatggcagaaattc<br>gaaagcaaattcgacccggtcgtcggttcagggcagggtcgttaaatagccgcttatgtctattgctggtttaccggtttattgactaccggaagcagtgtgaccgt<br>gtgcttctcaaatgcctgaggccagtttgctcaggctctccccgtggaggtaataattgctcgacatgaccaaaatcccttaacgtgagttttcgttccactgagcgt<br>cagaccccgtagaaaagatcaaaggatcttcttgagatcctttttttctgcgcgtaatctgctgcttgcaaacaaaaaaaccaccgctaccagcggtggtttgtttg<br>ccggatcaagagctaccaactctttttccgaaggtaactggcttcagcagagcgcagataccaaatactgtccttctagtgtagccgtagttaggccaccacttc<br>aagaactctgtagcaccgcctacatacctcgctctgctaatcctgttaccagtggctgctgccagtggcgataagtcgtgtcttaccgggttggactcaagacgat<br>agttaccggataaggcgcagcggtcgggctgaacggggggttcgtgcacacagcccagcttggagcgaacgacctacaccgaactgagatacctacagc<br>gtgagcattgagaaagcgccacgcttcccgaagggagaaaggcggacaggtatccggtaagcggcagggtcggaacaggagagcgcacgagggagc<br>ttccagggggaaacgcctggtatctttatagtcctgtcgggtttcgccacctctgacttgagcgtcgatttttgtgatgctcgtcaggggggccgagcctatggaaa<br>aacgccagcaacgcggcctttttacggttcctggccttttgctggccttttgctcacatgttctttcctgcgttatcccctgattctgtggataaccgtattaccgcctttga<br>gtgagctgataccgctcgccgcagccgaacgaccgagcgcagcgagtcagtgagcgaggaagcggaagagcgc |
| pCT anti-GFP Nick 90 | ccaatacgcaaaccgcctctccccgcgcgttggccgattcattaatgcagctggcacgacaggtttcccgactggaaagcgggcagtgagcgcaacgcaatt<br>aatgtgagttacctcactcattaggcaccccaggctttacactttatgcttccggctcctatgttgtgtggaattgtgagcggataacaatttcacacaggaaacag<br>ctatgaccatgattacgccaagctgccagatctgcagccgctatatatccgcggattaacgcgctagccggctgggccgtcgaacggaattaaccctcactaa<br>agggaacaaaagctgggtacccgacaggttatcagcaacaacacagtcatatccattctcaattagctctaccacagtgtgtgaaccaatgtatccagcacca<br>cctgtaaccaaaacaattttagaagtactttcactttgtaactgagctgtcatttatattgaattttcaaaaattcttactttttttttggatggacgcaaagaagtttaata<br>atcatattacatggcattaccaccatatacatatccatatacatatccatatctaatcttacttatatgttgtggaaatgtaaagagccccattatcttagcctaaaaaa<br>accttctctttggaactttcagtaatacgcttaactgctcattgctatattgaagtacggattagaagccgccgagcgggtgacagccctccgaaggaagactctc<br>ctccgtgcgtcctcgtcttcaccggtcgcgttcctgaaacgcagatgtgcctcgcgccgcactgctccgaacaataaagattctacaatactagcttttatggttatg<br>aagaggaaaaattggcagtaacctggccccacaaaccttcaaatgaacgaatcaaattaacaaccataggatgataatgcgattagtttttttagccttatttctg<br>gggtaattaatcagcgaagcgatgatttttgatctattaacagatatataaatgcaaaaactgcataaccactttaactaatactttcaacattttcggtttgtattactt<br>cttattcaaatgtaataaaagtatcaacaaaaaattgttaatatacctctatactttaacgtcaaggagaaaaaaccccggatcgaattccctacttcatacattttc<br>aattaagatgcagttacttcgctgtttttcaatattttctgttattgcttcagttttagcacaggaactgacaactatatgcgagcaaatcccctcaccaactttagaatc<br>gacgccgtactctttgtcaacgactactattttggccaacgggaaggcaatgcaaggagtttttgaatattacaaatcagtaacgtttgtcagtaattgcggttctca<br>cccctcaacaactagcaaaggcagccccataaacacacagtatgtttttaaggacaatagctcgacgattgaaggtagatacccatacgacgttccagacta<br>cgctctgcaggctagtggtggcggaggttctggtggaggcgggtctggtgggggaggatctgccatggtgcagctggtggaaagcggtggtgcactggtgca<br>gccaggtggcagcctgcgtctgagctgcgcggcgagcggctttccggtgaaccgctatagcatgcgctggtatcgtcaggcgccgggcaaagaacgcgaat<br>gggtggcgggcatgagcagcgcgggcgatcgcagcagctatgaagatagcgtgaaaggccgctttaccattagccgcgatgatgcgcgcaacaccgtgta<br>tctgcagatgaacagcctgaaaccggaagataccgcggtgtattattgcaacgtgaacgtgggctttgaatattggggccagggcacccaggtgaccgtgag<br>cagcgcggccgctagcgaaaccccaggtacgagcgaaagcgcgaccccggaacatatggtgccgaccatcgtgatggtggacgcctacaagcgttaca<br>agtaatagctcgagatctgataacaacagtgtagatgtaacaaaatcgactttgttcccactgtacttttagctcgtacaaaatacaatatacttttcatttctccgtaa<br>acaacatgttttcccatgtaatatcctttttctatttttcgttccgttaccaactttacacatactttatatagctattcacttctatacactaaaaaactaagacaattttaattt |

| | |
|---|---|
| | tgctgcctgccatatttcaatttgttataaattcctataatttatcctattagtagctaaaaaaagatgaatgtgaatcgaatcctaagagaattgag**CCTCAGCAGGGGCATCCCTCCTTTCAAGATAAATAATTTATACACTATTCTATTGGAATCTTAATCATTCTGGCCGATTCATTAATGCAGTTTGCTGAGG**ctccaattcgccctatagtgagtcgtattacaattcactggccgtcgttttacaacgtcgtgactgggaaaaccctggcgttacccaacttaatcgccttgcagcacatccccctttcgccagctggcgtaatagcgaagaggcccgcaccgatcgcctttcccaacagttgcggagcctgaatggcgaatggacgcgccctgtagcggcgcattaagcgcggcgggtgtggtggttacgcgcagcgtgaccgctacacttgccagcgccctagcgcccgctcctttcgctttcttcccttcctttctcgccacgttcgccggctttccccgtcaagctctaaatcgggggctccctttagggttccgatttagtgctttacggcacctcgaccccaaaaaacttgattagggtgatggttcacgtagtgggccatcgccctgatagacggtttttcgccctttgacgttggagtccacgttctttaatagtggactcttgttccaaactggaacaacactcaaccctatctcggtctattcttttgatttataagggattttgccgatttcggcctattggttaaaaaatgagctgatttaacaaaaatttaacgcgaattttaacaaaatattaacgcttacaatttcctgatgcggtattttctccttacgcatctgtgcggtatttcacaccgcatagatcggcaagtgcacaaacaatacttaaataaatactactcagtaataacctatttcttagcatttttgacgaaatttgctattttgttagagtcttttacaccatttgtctccacacctccgcttacatcaacaccaataacgccatttaatctaagcgcatcaccaacattttctggcgtcagtccaccagctaacataaaatgtaagctttcggggctctcttgccttccaacccagtcagaaatcgagttccaatccaaaagttcacctgtcccacctgcttctgaatcaaacaagggaataaacgaatgaggtttctgtgaagctgcactgagtagtatgttgcagtcttttggaaatacgagtcttttaataactggcaaaccgaggaactcttggtattcttgccacgactcatctccatgcagttggacgatatcaatgccgtaatcattgaccagagccaaaacatcctccttaggttgattacgaaacacgccaaccaagtatttcggagtgcctgaactatttttatatgcttttacaagacttgaaattttccttgcaataaccgggtcaattgttctctttctattgggcacacatataatacccagcaagtcagcatcggaatctagagcacattctgcggcctctgtgctctgcaagccgcaaactttcaccaatggaccagaactacctgtgaaattaataacagacatactccaagctgcctttgtgtgcttaatcacgtatactcacgtgctcaatagtcaccaatgccctccctcttggccctctccttttcttttttcgaccgaattaattcttaatcggcaaaaaaagaaaagctccggatcaagattgtacgtaaggtgacaagctatttttcaataaagaatatcttccactactgccatctggcgtcataactgcaaagtacacatatattacgatgctgtctattaaatgcttcctatattatatatatagtaatgtcgtttatggtgcactctcagtacaatctgctctgatgccgcatagttaagccagccccgacacccgccaacacccgctgacgcgccctgacgggcttgtctgctcccggcatccgcttacagacaagctgtgaccgtctccgggagctgcatgtgtcagaggttttcaccgtcatcaccgaaacgcgcgagacgaaagggcctcgtgatacgcctatttttataggttaatgtcatgataataatggtttcttaggacggatcgcttgcctgtaacttacacgcgcctcgtatcttttaatgatggaataatttgggaatttactctgtgtttatttatttttatgttttgtatttggattttagaaagtaaataaagaaggtagaagagttacggaatgaagaaaaaaaaataaacaaaggtttaaaaaatttcaacaaaaagcgtactttacatatatatttattagacaagaaaagcagattaaatagatatacattcgattaacgataagtaaaatgtaaaatcacaggattttcgtgtgtggtcttctacacagacaagatgaaacaattcggcattaatacctgagagcaggaagagcaagataaaaggtagtatttgttggcgatccccctagagtcttttacatcttcggaaaacaaaaactatttttctttaatttctttttttactttctatttttaatttatatatttatattaaaaaatttaaattataattatttttatagcacgtgatgaaaaggacccaggtggcacttttcggggaaatgtgcgcggaacccctatttgtttatttttctaaatacattcaaatatgtatccgctcatgtcgagacgttgggtgaggttccaactttcaccataatgaaataagatcactaccgggcgtattttttgagttatcgagattttcaggagctaaggaagctaaaatggagaaaaaaatcactggatataccaccgttgatatatcccaatggcatcgtaaagaacattttgaggcatttcagtcagttgctcaatgtacctataaccagaccgttcagctggatattacggcctttttaaagaccgtaaagaaaaataagcacaagttttatccggcctttattcacattcttgcccgcctgatgaatgctcacccggagttccgtatggcaatgaaagacggtgagctggtgatatgggatagtgttcacccttgttacaccgttttccatgagcaaactgaaacgttttcatcgctctggagtgaataccacgacgatttccggcagtttctacacatatattcgcaagatgtggcgtgttacggtgaaaacctggcctatttccctaaagggtttattgagaatatgtttttcgtctcagccaatccctgggtgagtttcaccagttttgatttaaacgtggctaatatggacaacttcttcgcccccgttttcacgatgggcaaatattatacgcaaggcgacaaggtgctgatgccgctggcgattcaggttcatcatgccgtttgtgatggcttccatgtcggcagaatgcttaatgaattacaacagtactgcgatgagtggcagggcggggcgtaatttttttaaggcagttattggtgcccttaaacgcctggtgctacgcctgaataagtgataataagcggatgaatggcagaaattcgaaagcaaattcgacccggtcgtcggttcagggcagggtcgttaaatagccgcttatgtctattgctggtttaccggtttattgactaccggaagcagtgtgaccgtgtgcttctcaaatgcctgaggccagtttgctcaggctctccccgtggaggtaataattgctcgacatgaccaaaatcccttaacgtgagttttcgttccactgagcgtcagaccccgtagaaaagatcaaaggatcttcttgagatcctttttttctgcgcgtaatctgctgcttgcaaacaaaaaaaccaccgctaccagcggtggtttgtttgccggatcaagagctaccaactctttttccgaaggtaactggcttcagcagagcgcagataccaaatactgtccttctagtgtagccgtagttaggccaccacttcaagaactctgtagcaccgcctacatacctcgctctgctaatcctgttaccagtggctgctgccagtggcgataagtcgtgtcttaccgggttggactcaagacgatagttaccggataaggcgcagcggtcgggctgaacggggggttcgtgcacacagcccagcttggagcgaacgacctacaccgaactgagatacctacagcgtgagcattgagaaagcgccacgcttcccgaagggagaaaggcggacaggtatccggtaagcggcagggtcggaacaggagagcgcacgagggagcttccagggggaaacgcctggtatctttatagtcctgtcgggtttcgccacctctgacttgagcgtcgatttttgtgatgctcgtcaggggggcggagcctatggaaaaacgccagcaacgcggcctttttacggttcctggccttttgctggccttttgctcacatgttctttcctgcgttatcccctgattctgtggataaccgtattaccgcctttgagtgagctgataccgctcgccgcagccgaacgaccgagcgcagcgagtcagtgagcgaggaagcggaagagcgc |

| | |
|---|---|
| pCT anti-GFP FspI 30 | cgcctctccccgcgcgttggccgattcattaatgcagctggcacgacaggtttcccgactggaaagcgggcagtgagcgcaacgcaattaatgtgagttacct<br>cactcattaggcaccccaggctttacactttatgcttccggctcctatgttgtgtggaattgtgagcggataacaatttcacacaggaaacagctatgaccatgatt<br>acgccaagctgccagatctgcagccgctatatatccgcggattaacgcgctagccggctgggccgtcgaacggaattaaccctcactaaagggaacaaaa<br>gctgggtacccgacaggttatcagcaacaacacagtcatatccattctcaattagctctaccacagtgtgtgaaccaatgtatccagcaccacctgtaaccaaa<br>acaattttagaagtactttcactttgtaactgagctgtcatttatattgaattttcaaaaattcttactttttttttggatggacgcaaagaagtttaataatcatattacatgg<br>cattaccaccatatacatatccatatacatatccatatctaatcttacttatatgttgtggaaatgtaaagagccccattatcttagcctaaaaaaaccttctctttgga<br>actttcagtaatacgcttaactgctcattgctatattgaagtacggattagaagccgccgagcgggtgacagccctccgaaggaagactctcctccgtgcgtcct<br>cgtcttcaccggtcgcgttcctgaaacgcagatgtgcctcgcgccgcactgctccgaacaataaagattctacaatactagcttttatggttatgaagaggaaaa<br>attggcagtaacctggccccacaaaccttcaaatgaacgaatcaaattaacaaccataggatgataatgcgattagttttttagccttatttctggggtaattaatc<br>agcgaagcgatgatttttgatctattaacagatatataaatgcaaaaactgcataaccactttaactaatactttcaacattttcggtttgtattacttcttattcaaatgt<br>aataaaagtatcaacaaaaaattgttaatatacctctatactttaacgtcaaggagaaaaaaccccggatcgaattccctacttcatacattttcaattaagatgc<br>agttacttcgctgtttttcaatattttctgttattgcttcagttttagcacaggaactgacaactatatgcgagcaaatcccctcaccaactttagaatcgacgccgtact<br>ctttgtcaacgactactattttggccaacgggaaggcaatgcaaggagtttttgaatattacaaatcagtaacgtttgtcagtaattgcggttctcacccctcaacaa<br>ctagcaaaggcagccccataaacacacagtatgtttttaaggacaatagctcgacgattgaaggtagatacccatacgacgttccagactacgctctgcaggc<br>tagtggtggcggaggttctggtggaggcgggtctggtgggggaggatctgccatggtgcagctggtggaaagcggtggtgcactggtgcagccaggtggca<br>gcctgcgtctgagctgcgcggcgagcggctttccggtgaaccgctatagcatgcgctggtatcgtcaggcgccgggcaaagaacgcgaatgggtggcgggc<br>atgagcagcgcgggcgatcgcagcagctatgaagatagcgtgaaaggccgctttaccattagccgcgatgatgcgcgcaacaccgtgtatctgcagatgaa<br>cagcctgaaaccggaagataccgcggtgtattattgcaacgtgaacgtgggctttgaatattggggccagggcacccaggtgaccgtgagcagcgcggccg<br>ctagcgaaaccccaggtacgagcgaaagcgcgaccccggaacatatggtgccgaccatcgtgatggtggacgcctacaagcgttacaagtaatagctcga<br>gatctgataacaacagtgtagatgtaacaaaatcgactttgttcccactgtacttttagctcgtacaaaatacaatatacttttcatttctccgtaaacaacatgttttcc<br>catgtaatatccttttctatttttcgttccgttaccaactttacacatactttatatagctattcacttctatacactaaaaaactaagacaattttaattttgctgcctgccat<br>atttcaatttgttataaattcctataatttatcctattagtagctaaaaaaagatgaatgtgaatcgaatcctaagagaattgag**CCTCAGCAGGTGGCC**<br>**GATTCATTAATGCAGTTTGCGCAGGTGGCCGATTCATTAATGCAGTTTGCTGAGG**ctccaattcgccctatagtgagtc<br>gtattacaattcactggccgtcgttttacaacgtcgtgactgggaaaaccctggcgttacccaacttaatcgccttgcagcacatcccccttcgccagctggcgta<br>atagcgaagaggcccgcaccgatcgcccttcccaacagttgggcagcctgaatggcgaatggacgcgccctgtagcggcgcattaagcgcggcgggtgtg<br>gtggttacgcgcagcgtgaccgctacacttgccagcgccctagcgcccgctcctttcgctttcttcccttcctttctcgccacgttcgccggctttccccgtcaagctc<br>taaatcgggggctccctttagggttccgatttagtgctttacggcacctcgaccccaaaaaacttgattagggtgatggttcacgtagtgggccatcgccctgata<br>gacggtttttcgccctttgacgttggagtccacgttctttaatagtggactcttgttccaaactggaacaacactcaaccctatctcggtctattcttttgatttataaggg<br>attttgccgatttcggcctattggttaaaaaatgagctgatttaacaaaaatttaacgcgaattttaacaaaatattaacgcttacaatttcctgatgcggtattttctcct<br>tacgcatctgtgcggtatttcacaccgcatagatcggcaagtgcacaaacaatacttaaataaatactactcagtaataacctatttcttagcatttttgacgaaattt<br>gctattttgttagagtcttttacaccatttgtctccacacctccgcttacatcaacaccaataacgccatttaatctaagcgcatcaccaacattttctggcgtcagtcc<br>accagctaacataaaatgtaagctttcggggctctcttgccttccaacccagtcagaaatcgagttccaatccaaaagttcacctgtcccacctgcttctgaatca<br>aacaagggaataaacgaatgaggtttctgtgaagctgcactgagtagtatgttgcagtcttttggaaatacgagtcttttaataactggcaaaccgaggaactctt<br>ggtattcttgccacgactcatctccatgcagttggacgatatcaatgccgtaatcattgaccagagccaaaacatcctccttaggttgattacgaaacacgccaa<br>ccaagtatttcggagtgcctgaactatttttatatgcttttacaagacttgaaattttccttgcaataaccgggtcaattgttctctttctattgggcacacatataataccc<br>agcaagtcagcatcggaatctagagcacattctgcggcctctgtgctctgcaagccgcaaactttcaccaatggaccagaactacctgtgaaattaataacag<br>acatactccaagctgcctttgtgtgcttaatcacgtatactcacgtgctcaatagtcaccaatgccctccctcttggccctctccttttcttttttcgaccgaattaattctt<br>aatcggcaaaaaaagaaaagctccggatcaagattgtacgtaaggtgacaagctatttttcaataaagaatatcttccactactgccatctggcgtcataactgc<br>aaagtacacatatattacgatgctgtctattaaatgcttcctatattatatatatagtaatgtcgtttatggtgcactctcagtacaatctgctctgatgccgcatagttaa<br>gccagccccgacacccgccaacacccgctgacgcgccctgacgggcttgtctgctcccggcatccgcttacagacaagctgtgaccgtctccgggagctgc<br>atgtgtcagaggttttcaccgtcatcaccgaaacgcgcgagacgaaagggcctcgtgatacgcctatttttataggttaatgtcatgataataatggtttcttagga<br>cggatcgcttgcctgtaacttacacgcgcctcgtatcttttaatgatggaataatttgggaatttactctgtgtttatttatttttatgttttgtatttggattttagaaagtaaat<br>aaagaaggtagaagagttacggaatgaagaaaaaaaaaataaacaaaggtttaaaaaatttcaacaaaaagcgtactttacatatatatttattagacaagaa<br>aagcagattaaatagatatacattcgattaacgataagtaaaatgtaaaatcacaggattttcgtgtgtggtcttctacacagacaagatgaaacaattcggcatt |

| | |
|---|---|
| | aatacctgagagcaggaagagcaagataaaaggtagtatttgttggcgatcccctagagtcttttacatcttcggaaaacaaaaactattttttctttaatttcttttttactttctatttttaatttatatatttatattaaaaaatttaaattataattatttttatagcacgtgatgaaaaggacccaggtggcacttttcggggaaatgtgcgcggaaccccctatttgtttatttttctaaatacattcaaatatgtatccgctcatgtcgagacgttgggtgaggttccaactttcaccataatgaaataagatcactaccgggcgtattttttgagttatcgagattttcaggagctaaggaagctaaaatggagaaaaaaatcactggatataccaccgttgatatatcccaatggcatcgtaaagaacattttgaggcatttcagtcagttgctcaatgtacctataaccagaccgttcagctggatattacggcctttttaaagaccgtaaagaaaaataagcacaagttttatccggcctttattcacattcttgcccgcctgatgaatgctcacccggagttccgtatggcaatgaaagacggtgagctggtgatatgggatagtgttcacccttgttacaccgttttccatgagcaaactgaaacgttttcatcgctctggagtgaataccacgacgatttccggcagtttctacacatatattcgcaagatgtggcgtgttacggtgaaaacctggcctatttccctaaagggtttattgagaatatgtttttcgtctcagccaatccctgggtgagtttcaccagttttgatttaaacgtggctaatatggacaacttcttcgcccccgttttcacgatgggcaaatattatacgcaaggcgacaaggtgctgatgccgctggcgattcaggttcatcatgccgtttgtgatggcttccatgtcggcagaatgcttaatgaattacaacagtactgcgatgagtggcagggcggggcgtaatttttttaaggcagttattggtgcccttaaacgcctggtgctacgcctgaataagtgataataagcggatgaatggcagaaattcgaaagcaaattcgacccggtcgtcggttcagggcagggtcgttaaatagccgcttatgtctattgctggtttaccggtttattgactaccggaagcagtgtgaccgtgtgcttctcaaatgcctgaggccagtttgctcaggctctccccgtggaggtaataattgctcgacatgaccaaaatcccttaacgtgagttttcgttccactgagcgtcagaccccgtagaaaagatcaaaggatcttcttgagatcctttttttctgcgcgtaatctgctgcttgcaaacaaaaaaaccaccgctaccagcggtggtttgtttgccggatcaagagctaccaactctttttccgaaggtaactggcttcagcagagcgcagataccaaatactgtccttctagtgtagccgtagttaggccaccacttcaagaactctgtagcaccgcctacatacctcgctctgctaatcctgttaccagtggctgctgccagtggcgataagtcgtgtcttaccgggttggactcaagacgatagttaccggataaggcgcagcggtcgggctgaacggggggttcgtgcacacagcccagcttggagcgaacgacctacaccgaactgagatacctacagcgtgagcattgagaaagcgccacgcttcccgaagggagaaaggcggacaggtatccggtaagcggcagggtcggaacaggagagcgcacgagggagcttccagggggaaacgcctggtatctttatagtcctgtcgggtttcgccacctctgacttgagcgtcgatttttgtgatgctcgtcaggggggcggagcctatggaaaaacgccagcaacgcggcctttttacggttcctggccttttgctggccttttgctcacatgttctttcctgcgttatcccctgattctgtggataaccgtattaccgcctttgagtgagctgataccgctcgccgcagccgaacgaccgagcgcagcgagtcagtgagcgaggaagcggaagagcgcccaatacgcaaac |
| pCT anti-GFP FspI 90 | ccaatacgcaaaccgcctctccccgcgcgttggccgattcattaatgcagctggcacgacaggtttcccgactggaaagcgggcagtgagcgcaacgcaattaatgtgagttacctcactcattaggcaccccaggctttacactttatgcttccggctcctatgttgtgtggaattgtgagcggataacaatttcacacaggaaacagctatgaccatgattacgccaagctgccagatctgcagccgctatatatccgcggattaacgcgctagccggctgggccgtcgaacggaattaaccctcactaaagggaacaaaagctgggtacccgacaggttatcagcaacaacacagtcatatccattctcaattagctctaccacagtgtgtgaaccaatgtatccagcaccacctgtaaccaaaacaattttagaagtactttcactttgtaactgagctgtcatttatattgaattttcaaaaattcttactttttttttggatggacgcaaagaagtttaataatcatattacatggcattaccaccatatacatatccatatacatatccatatctaatcttacttatatgttgtggaaatgtaaagagccccattatcttagcctaaaaaaaccttctctttggaactttcagtaatacgcttaactgctcattgctatattgaagtacggattagaagccgccgagcgggtgacagccctccgaaggaagactctcctccgtgcgtcctcgtcttcaccggtcgcgttcctgaaacgcagatgtgcctcgcgccgcactgctccgaacaataaagattctacaatactagcttttatggttatgaagaggaaaaattggcagtaacctggccccacaaaccttcaaatgaacgaatcaaattaacaaccataggatgataatgcgattagttttttagccttatttctggggtaattaatcagcgaagcgatgatttttgatctattaacagatatataaatgcaaaaactgcataaccactttaactaatactttcaacattttcggtttgtattacttcttattcaaatgtaataaaagtatcaacaaaaaattgttaatatacctctatactttaacgtcaaggagaaaaaaccccggatcgaattccctacttcatacattttcaattaagatgcagttacttcgctgtttttcaatattttctgttattgcttcagttttagcacaggaactgacaactatatgcgagcaaatcccctcaccaactttagaatcgacgccgtactctttgtcaacgactactattttggccaacgggaaggcaatgcaaggagtttttgaatattacaaatcagtaacgtttgtcagtaattgcggttctcacccctcaacaactagcaaaggcagccccataaacacacagtatgtttttaaggacaatagctcgacgattgaaggtagatacccatacgacgttccagactacgctctgcaggctagtggtggcggaggttctggtggaggcgggtctggtgggggaggatctgccatggtgcagctggtggaaagcggtggtgcactggtgcagccaggtggcagcctgcgtctgagctgcgcggcgagcggctttccggtgaaccgctatagcatgcgctggtatcgtcaggcgccgggcaaagaacgcgaatgggtggcgggcatgagcagcgcgggcgatcgcagcagctatgaagatagcgtgaaaggccgctttaccattagccgcgatgatgcgcgcaacaccgtgtatctgcagatgaacagcctgaaaccggaagataccgcggtgtattattgcaacgtgaacgtgggctttgaatattggggccagggcacccaggtgaccgtgagcagcgcggccgctagcgaaaccccaggtacgagcgaaagcgcgaccccggaacatatggtgccgaccatcgtgatggtggacgcctacaagcgttacaagtaatagctcgagatctgataacaacagtgtagatgtaacaaaatcgactttgttcccactgtacttttagctcgtacaaaatacaatatacttttcatttctccgtaaacaacatgttttcccatgtaatatccttttctatttttcgttccgttaccaactttacacatactttatatagctattcacttctatacactaaaaaactaagacaattttaatttgctgcctgccatatttcaatttgttataaattcctataatttatcctattagtagctaaaaaaagatgaatgtgaatcgaatcctaagagaattgag**CCTCAGCAGGGGCATCCCTCCTTTCAAGATAAATAATTTATACACTATTCTATTGGAATCTTAATCATTCTGGCCGATT** |

**CATTAATGCAGTTTGCGCAGGGGCATCCCTCCTTTCAAGATAAATAATTTATACACTATTCTATTGGAATCT**
**TAATCATTCTGGCCGATTCATTAATGCAGTTTGCTGAGG**ctccaattcgccctatagtgagtcgtattacaattcactggccgtcgtttt
acaacgtcgtgactgggaaaaccctggcgttacccaacttaatcgccttgcagcacatccccctttcgccagctggcgtaatagcgaagaggcccgcaccga
tcgccttcccaacagttgcgcagcctgaatggcgaatggacgcgccctgtagcggcgcattaagcgcggcgggtgtggtggttacgcgcagcgtgaccgct
acacttgccagcgccctagcgcccgctcctttcgctttcttcccttcctttctcgccacgttcgccggctttccccgtcaagctctaaatcgggggctccctttagggtt
ccgatttagtgctttacggcacctcgaccccaaaaaacttgattagggtgatggttcacgtagtgggccatcgccctgatagacggtttttcgccctttgacgttgga
gtccacgttctttaatagtggactcttgttccaaactggaacaacactcaaccctatctcggtctattcttttgatttataagggattttgccgatttcggcctattggttaa
aaaatgagctgatttaacaaaaatttaacgcgaattttaacaaaatattaacgcttacaatttcctgatgcggtattttctccttacgcatctgtgcggtatttcacacc
gcatagatcggcaagtgcacaaacaatacttaaataaatactactcagtaataacctatttcttagcatttttgacgaaatttgctattttgttagagtcttttacaccat
ttgtctccacacctccgcttacatcaacaccaataacgccatttaatctaagcgcatcaccaacattttctggcgtcagtccaccagctaacataaaatgtaagctt
tcggggctctcttgccttccaacccagtcagaaatcgagttccaatccaaaagttcacctgtcccacctgcttctgaatcaaacaagggaataaacgaatgagg
tttctgtgaagctgcactgagtagtatgttgcagtcttttggaaatacgagtcttttaataactggcaaaccgaggaactcttggtattcttgccacgactcatctccat
gcagttggacgatatcaatgccgtaatcattgaccagagccaaaacatcctccttaggttgattacgaaacacgccaaccaagtatttcggagtgcctgaacta
tttttatatgcttttacaagacttgaaattttccttgcaataaccgggtcaattgttctctttctattgggcacacatataatacccagcaagtcagcatcggaatctaga
gcacattctgcggcctctgtgctctgcaagccgcaaactttcaccaatggaccagaactacctgtgaaattaataacagacatactccaagctgcctttgtgtgct
taatcacgtatactcacgtgctcaatagtcaccaatgccctccctcttggccctctccttttcttttttcgaccgaattaattcttaatcggcaaaaaaagaaaagctcc
ggatcaagattgtacgtaaggtgacaagctatttttcaataaagaatatcttccactactgccatctggcgtcataactgcaaagtacacatatattacgatgctgtc
tattaaatgcttcctatattatatatatagtaatgtcgtttatggtgcactctcagtacaatctgctctgatgccgcatagttaagccagccccgacacccgccaacac
ccgctgacgcgccctgacgggcttgtctgctcccggcatccgcttacagacaagctgtgaccgtctccgggagctgcatgtgtcagaggttttcaccgtcatcac
cgaaacgcgcgagacgaaagggcctcgtgatacgcctatttttataggttaatgtcatgataataatggtttcttaggacggatcgcttgcctgtaacttacacgcg
cctcgtatcttttaatgatggaataatttgggaatttactctgtgtttattatttttatgttttgtatttggattttagaaagtaaataaagaaggtagaagagttacggaatg
aagaaaaaaaaataaacaaaggtttaaaaaatttcaacaaaaagcgtactttacatatatatttattagacaagaaaagcagattaaatagatatacattcgat
taacgataagtaaaatgtaaaatcacaggattttcgtgtgtggtcttctacacagacaagatgaaacaattcggcattaatacctgagagcaggaagagcaag
ataaaaggtagtatttgttggcgatccccctagagtcttttacatcttcggaaaacaaaaactattttttctttaatttctttttttactttctatttttaatttatatatttatattaa
aaaatttaaattataattatttttatagcacgtgatgaaaaggacccaggtggcacttttcggggaaatgtgcgcggaacccctatttgtttatttttctaaatacattca
aatatgtatccgctcatgtcgagacgttgggtgaggttccaactttcaccataatgaaataagatcactaccgggcgtattttttgagttatcgagattttcaggagct
aaggaagctaaaatggagaaaaaaatcactggatataccaccgttgatatatcccaatggcatcgtaaagaacattttgaggcatttcagtcagttgctcaatgt
acctataaccagaccgttcagctggatattacggcctttttaaagaccgtaaagaaaaataagcacaagttttatccggcctttattcacattcttgcccgcctgatg
aatgctcacccggagttccgtatggcaatgaaagacggtgagctggtgatatgggatagtgttcacccttgttacaccgttttccatgagcaaactgaaacgttttc
atcgctctggagtgaataccacgacgatttccggcagtttctacacatatattcgcaagatgtggcgtgttacggtgaaaacctggcctatttccctaaagggtttat
tgagaatatgtttttcgtctcagccaatccctgggtgagtttcaccagttttgatttaaacgtggctaatatggacaacttcttcgcccccgttttcacgatgggcaaat
attatacgcaaggcgacaaggtgctgatgccgctggcgattcaggttcatcatgccgtttgtgatggcttccatgtcggcagaatgcttaatgaattacaacagta
ctgcgatgagtggcagggcggggcgtaatttttttaaggcagttattggtgcccttaaacgcctggtgctacgcctgaataagtgataataagcggatgaatggc
agaaattcgaaagcaaattcgacccggtcgtcggttcagggcagggtcgttaaatagccgcttatgtctattgctggtttaccggtttattgactaccggaagcagt
gtgaccgtgtgcttctcaaatgcctgaggccagtttgctcaggctctccccgtggaggtaataattgctcgacatgaccaaaatcccttaacgtgagttttcgttcca
ctgagcgtcagaccccgtagaaaagatcaaaggatcttcttgagatcctttttttctgcgcgtaatctgctgcttgcaaacaaaaaaaccaccgctaccagcggt
ggtttgtttgccggatcaagagctaccaactctttttccgaaggtaactggcttcagcagagcgcagataccaaatactgtccttctagtgtagccgtagttaggcc
accacttcaagaactctgtagcaccgcctacatacctcgctctgctaatcctgttaccagtggctgctgccagtggcgataagtcgtgtcttaccgggttggactca
agacgatagttaccggataaggcgcagcggtcgggctgaacggggggttcgtgcacacagcccagcttggagcgaacgacctacaccgaactgagatac
ctacagcgtgagcattgagaaagcgccacgcttcccgaagggagaaaggcggacaggtatccggtaagcggcagggtcggaacaggagagcgcacga
gggagcttccagggggaaacgcctggtatctttatagtcctgtcgggtttcgccacctctgacttgagcgtcgatttttgtgatgctcgtcaggggggcgagccta
tggaaaaacgccagcaacgcggcctttttacggttcctggccttttgctggccttttgctcacatgttctttcctgcgttatcccctgattctgtggataaccgtattaccg
cctttgagtgagctgataccgctcgccgcagccgaacgaccgagcgcagcgagtcagtgagcgaggaagcggaagagcgc

| | |
|---|---|
| PF<br>Nbb102<br>CAM<br>Nick 10 | tggccttttgctcacatgcgcccaatacgcaaaccgcctctccccgcgcgttggccgattcattaatgcaggaatgattaagattccaatagaatagtgtataaatt<br>atttatcttgaaaggagggatgccctggcacgac**CCTCAGCAGGTTTGCTGAGG**cccgactggaaagcgggcagtgagcgcaacgcaatta<br>atgtgagttagctcactcattaggcaccccaggctttacactttatgcttccggctcgtatgttgtgtggaattgtgagcggataacaatttgaattcaaggagacag<br>tcataatgaaatacctattgcctacggcggccgctggattgttattactcgcggcccagccggcaatggcacaggtccagttacaagagtcaggcggggggctt<br>gtccaggccggggggtcactgcgtttatcgtgtgcggcaagtggatacatcagcgacgcttactacatgggatggtatcgccaggcccctgggaaagaacgc<br>gaatttgtggctaccattacccacgggactaacacttactacgcggattccgtaaaagggcgcttcaccatcagccgcgataacgcaaagaacactgtatatct<br>gcaaatgaatagcttaaagcctgaagataccgcggtctattactgtgccgtacttgaaacacgttcttattctttccgctattggggccagggaactcaggtcactg<br>tatcgagccaccatcaccaccatcatggcgcagaacaaaaactcatctcagaagaggatctgtcttaggccgaaactgttgaaagttgtttagcaaaacctcat<br>acagaaaattcatttactaacgtctggaaagacgacaaaactttagatcgttacgctaactatgagggctgtctgtggaatgctacaggcgttgtggtttgtactgg<br>tgacgaaactcagtgttacggtacatgggttcctattgggcttgctatccctgaaaatgagggtggtggctctgagggtggcggttctgagggtggcggttctgag<br>ggtggcggtactaaacctcctgagtacggtgatacacctattccgggctatacttatatcaaccctctcgacggcacttatccgcctggtactgagcaaaacccc<br>gctaatcctaatccttctcttgaggagtctcagcctcttaatactttcatgtttcagaataataggttccgaaataggcagggtgcattaactgtttatacgggcactgtt<br>actcaaggcactgaccccgttaaaacttattaccagtacactcctgtatcatcaaaagccatgtatgacgcttactggaacggtaaattcagagactgcgctttcc<br>attctggctttaatgaggatccattcgtttgtgaatatcaaggccaatcgtctgacctgcctcaacctcctgtcaatgctggcggcggctctggtggtggttctggtgg<br>cggctctgagggtggcggctctgagggtggcggttctgagggtggcggctctgagggtggcggttccggtggcggctccggttccggtgattttgattatgaaaa<br>aatggcaaacgctaataagggggctatgaccgaaaatgccgatgaaaacgcgctacagtctgacgctaaaggcaaacttgattctgtcgctactgattacggt<br>gctgctatcgatggtttcattggtgacgtttccggccttgctaatggtaatggtgctactggtgattttgctggctctaattcccaaatggctcaagtcggtgacggtgat<br>aattcacctttaatgaataatttccgtcaatatttaccttctttgcctcagtcggttgaatgtcgcccttatgtctttggcgctggtaaaccatatgaattttctattgattgtg<br>acaaaataaacttattccgtggtgtctttgcgtttcttttatatgttgccacctttatgtatgtattttcgacgtttgctaacatactgcgtaataaggagtcttaagctagct<br>aacagtcctatgaatcaactacttagatggtattagtgacctgtaacagagcattagcgcaaggtgatttttgacttcttgcgctaatttttgtcatcaaacctgtcgc<br>actccttaatattttgttaaaattcgcgttaaatttttgttaaatcagctcattttttaaccaataggccgaaatcggcaaaatcccttataaatcaaaagaatagaccg<br>agatagggttgagtgttgttccagtttggaacaagagtccactattaaagaacgtggactccaacgtcaaagggcgaaaaaccgtctatcagggcgatggcc<br>cactacgtgaaccatcaccctaatcaagttttttggggtcgaggtgccgtaaagcactaaatcggaaccctaaagggagcccccgatttagagcttgacgggg<br>aaagccggcgaacgtggcgagaaaggaagggaagaaagcgaaaggagcgggcgctagggcgctggcaagtgtagcggtcacgctgcgcgtaacca<br>ccacacccgccgcgcttaatgcgccgctacagggcgcgtcaggtggcacttttcggggaaatgtgcgcggaacccctatttgtttatttttctaaatacattcaaat<br>atgtatccgctcatgtcgagacgttgggtgaggttccaactttcaccataatgaaataagatcactaccgggcgtattttttgagttatcgagattttcaggagctaa<br>ggaagctaaaatggagaaaaaaatcactggatataccaccgttgatatatcccaatggcatcgtaaagaacattttgaggcatttcagtcagttgctcaatgtac<br>ctataaccagaccgttcagctggatattacggcctttttaaagaccgtaaagaaaaataagcacaagttttatccggcctttattcacattcttgcccgcctgatga<br>atgctcacccggagttccgtatggcaatgaaagacggtgagctggtgatatgggatagtgttcacccttgttacaccgttttccatgagcaaactgaaacgttttca<br>tcgctctggagtgaataccacgacgatttccggcagtttctacacatatattcgcaagatgtggcgtgttacggtgaaaacctggcctatttccctaaagggtttatt<br>gagaatatgtttttcgtctcagccaatccctgggtgagtttcaccagttttgatttaaacgtggctaatatggacaacttcttcgcccccgttttcaccatgggcaaata<br>ttatacgcaaggcgacaaggtgctgatgccgctggcgattcaggttcatcatgccgtttgtgatggcttccatgtcggcagaatgcttaatgaattacaacagtact<br>gcgatgagtggcagggcggggcgtaatttttttaaggcagttattggtgcccttaaacgcctggtgctacgcctgaataagtgataataagcggatgaatggcag<br>aaattcgaaagcaaattcgacccggtcgtcggttcagggcagggtcgttaaatagccgcttatgtctattgctggtttaccggtttattgactaccggaagcagtgt<br>gaccgtgtgcttctcaaatgcctgaggccagtttgctcaggctctccccgtggaggtaataattgctcgacatgaccaaaatcccttaacgtgagttttcgttccact<br>gagcgtcagaccccgtagaaaagatcaaaggatcttcttgagatcctttttttctgcgcgtaatctgctgcttgcaaacaaaaaaaccaccgctaccagcggtgg<br>tttgtttgccggatcaagagctaccaactctttttccgaaggtaactggcttcagcagagcgcagataccaaatactgttcttctagtgtagccgtagttaggccacc<br>acttcaagaactctgtagcaccgcctacatacctcgctctgctaatcctgttaccagtggctgctgccagtggcgataagtcgtgtcttaccgggttggactcaag<br>acgatagttaccggataaggcgcagcggtcgggctgaacggggggttcgtgcacacagcccagcttggagcgaacgacctacaccgaactgagatacctaa<br>cagcgtgagctatgagaaagcgccacgcttcccgaagggagaaaggcggacaggtatccggtaagcggcagggtcggaacaggagagcgcacgagg<br>gagcttccagggggaaacgcctggtatctttatagtcctgtcgggtttcgccacctctgacttgagcgtcgatttttgtgatgctcgtcaggggggcggagcctatg<br>gaaaaacgccagcaacgcggcctttttacggttcctggccttttgc |

| | |
|---|---|
| PF<br>Nbb102<br>Nick 15 | tggccttttgctcacatgcgcccaatacgcaaaccgcctctccccgcgcgttggccgattcattaatgcaggaatgattaagattccaatagaatagtgtataaatt<br>atttatcttgaaaggagggatgccctgg**CCTCAGCACGACAGGTTTGCTGAGG**cccgactggaaagcgggcagtgagcgcaacgcaatt<br>aatgtgagttagctcactcattaggcaccccaggctttacactttatgcttccggctcgtatgttgtgtggaattgtgagcggataacaatttgaattcaaggagaca<br>gtcataatgaaatacctattgcctacggcggccgctggattgttattactcgcggcccagccggcaatggcacaggtccagttacaagagtcaggcgggggc<br>ttgtccaggccgggggtcactgcgtttatcgtgtgcggcaagtggatacatcagcgacgcttactacatgggatggtatcgccaggcccctgggaaagaacg<br>cgaatttgtggctaccattacccacgggactaacacttactacgcggattccgtaaaagggcgcttcaccatcagccgcgataacgcaaagaacactgtatat<br>ctgcaaatgaatagcttaaagcctgaagataccgcggtctattactgtgccgtacttgaaacacgttcttattctttccgctattggggccagggaactcaggtcac<br>tgtatcgagccaccatcaccaccatcatggcgcagaacaaaaactcatctcagaagaggatctgtcttaggccgaaactgttgaaagttgtttagcaaaacctc<br>atacagaaaattcatttactaacgtctggaaagacgacaaaactttagatcgttacgctaactatgagggctgtctgtggaatgctacaggcgttgtggtttgtact<br>ggtgacgaaactcagtgttacggtacatgggttcctattgggcttgctatccctgaaaatgagggtggtggctctgagggtggcggttctgagggtggcggttctg<br>agggtggcggtactaaacctcctgagtacggtgatacacctattccgggctatacttatatcaaccctctcgacggcacttatccgcctggtactgagcaaaacc<br>ccgctaatcctaatccttctcttgaggagtctcagcctcttaatactttcatgtttcagaataataggttccgaaataggcagggtgcattaactgtttatacgggcact<br>gttactcaaggcactgaccccgttaaaacttattaccagtacactcctgtatcatcaaaagccatgtatgacgcttactggaacggtaaattcagagactgcgcttt<br>ccattctggctttaatgaggatccattcgtttgtgaatatcaaggccaatcgtctgacctgcctcaacctcctgtcaatgctggcggcggctctggtggtggttctggt<br>ggcggctctgagggtggcggctctgagggtggcggttctgagggtggcggctctgagggtggcggttccggtggcggctccggttccggtgattttgattatgaa<br>aaaatggcaaacgctaataagggggctatgaccgaaaatgccgatgaaaacgcgctacagtctgacgctaaaggcaaacttgattctgtcgctactgattac<br>ggtgctgctatcgatggtttcattggtgacgtttccggccttgctaatggtaatggtgctactggtgattttgctggctctaattcccaaatggctcaagtcggtgacggt<br>gataattcacctttaatgaataatttccgtcaatatttaccttctttgcctcagtcggttgaatgtcgcccttatgtctttggcgctggtaaaccatatgaattttctattgatt<br>gtgacaaaataaacttattccgtggtgtctttgcgtttcttttatatgttgccacctttatgtatgtattttcgacgtttgctaacatactgcgtaataaggagtcttaagcta<br>gctaacagtcctatgaatcaactacttagatggtattagtgacctgtaacagagcattagcgcaaggtgattttgacttcttgcgctaatttttgtcatcaaacctgtc<br>gcactccttaatattttgttaaaattcgcgttaaattttgttaaatcagctcattttttaaccaataggccgaaatcggcaaaatcccttataaatcaaaagaatagac<br>cgagatagggttgagtgttgttccagtttggaacaagagtccactattaaagaacgtggactccaacgtcaaagggcgaaaaaccgtctatcagggcgatgg<br>cccactacgtgaaccatcaccctaatcaagttttttggggtcgaggtgccgtaaagcactaaatcggaaccctaaagggagcccccgatttagagcttgacgg<br>ggaaagccggcgaacgtggcgagaaaggaagggaagaaagcgaaaggagcgggcgctagggcgctggcaagtgtagcggtcacgctgcgcgtaac<br>caccacacccgccgcgcttaatgcgccgctacagggcgcgtcaggtggcacttttcggggaaatgtgcgcggaacccctatttgtttatttttctaaatacattca<br>aatatgtatccgctcatgtcgagacgttgggtgaggttccaactttcaccataatgaaataagatcactaccgggcgtattttttgagttatcgagattttcaggagct<br>aaggaagctaaaatggagaaaaaaatcactggatataccaccgttgatatatcccaatggcatcgtaaagaacattttgaggcatttcagtcagttgctcaatgt<br>acctataaccagaccgttcagctggatattacggcctttttaaagaccgtaaagaaaaataagcacaagttttatccggcctttattcacattcttgcccgcctgatg<br>aatgctcacccggagttccgtatggcaatgaaagacggtgagctggtgatatgggatagtgttcacccttgttacaccgttttccatgagcaaactgaaacgttttc<br>atcgctctggagtgaataccacgacgatttccggcagtttctacacatatattcgcaagatgtggcgtgttacggtgaaaacctggcctatttccctaaagggtttat<br>tgagaatatgtttttcgtctcagccaatccctgggtgagtttcaccagttttgatttaaacgtggctaatatggacaacttcttcgcccccgttttcaccatgggcaaat<br>attatacgcaaggcgacaaggtgctgatgccgctggcgattcaggttcatcatgccgtttgtgatggcttccatgtcggcagaatgcttaatgaattacaacagta<br>ctgcgatgagtggcagggcggggcgtaatttttttaaggcagttattggtgcccttaaacgcctggtgctacgcctgaataagtgataataagcggatgaatggc<br>agaaattcgaaagcaaattcgacccggtcgtcggttcagggcagggtcgttaaatagccgcttatgtctattgctggtttaccggtttattgactaccggaagcagt<br>gtgaccgtgtgcttctcaaatgcctgaggccagtttgctcaggctctccccgtggaggtaataattgctcgacatgaccaaaatcccttaacgtgagttttcgttcca<br>ctgagcgtcagaccccgtagaaaagatcaaaggatcttcttgagatcctttttttctgcgcgtaatctgctgcttgcaaacaaaaaaaccaccgctaccagcggt<br>ggtttgtttgccggatcaagagctaccaactctttttccgaaggtaactggcttcagcagagcgcagataccaaatactgttcttctagtgtagccgtagttaggcc<br>accacttcaagaactctgtagcaccgcctacatacctcgctctgctaatcctgttaccagtggctgctgccagtggcgataagtcgtgtcttaccgggttggactca<br>agacgatagttaccggataaggcgcagcggtcgggctgaacggggggttcgtgcacacagcccagcttggagcgaacgacctacaccgaactgagatac<br>ctacagcgtgagctatgagaaagcgccacgcttcccgaagggagaaaggcggacaggtatccggtaagcggcagggtcggaacaggagagcgcacga<br>gggagcttccagggggaaacgcctggtatctttatagtcctgtcgggtttcgccacctctgacttgagcgtcgatttttgtgatgctcgtcaggggggcggagccta<br>tggaaaaacgccagcaacgcggcctttttacggttcctggccttttgc |
